# Integration of anatomy ontologies and evo-devo using structured Markov models suggests a new framework for modeling discrete phenotypic traits

**DOI:** 10.1101/188672

**Authors:** Sergei Tarasov

## Abstract

Modeling discrete phenotypic traits for either ancestral character state reconstruction or morphology-based phylogenetic inference suffers from ambiguities of character coding, homology assessment, dependencies, and selection of adequate models. These drawbacks occur because trait evolution is driven by two key processes – hierarchical and hidden – which are not accommodated simultaneously by the available phylogenetic methods. The hierarchical process refers to the dependencies between anatomical body parts, while the hidden process refers to the evolution of gene regulatory networks underlying trait development. Herein, I demonstrate that these processes can be efficiently modeled using structured Markov models equipped with hidden states, which resolves the majority of the problems associated with discrete traits. Integration of structured Markov models with anatomy ontologies can adequately incorporate the hierarchical dependencies, while the use of the hidden states accommodates hidden evolution of gene regulatory networks and substitution rate heterogeneity. I assess the new models using simulations and theoretical synthesis. The new approach solves the long-standing tail color problem (that aims at coding tail when it is absent) and presents a previously unknown issue called the “two-scientist paradox”. The latter issue refers to the confounding nature of the coding of a trait and the hidden processes driving the trait’s evolution; failing to account for the hidden process may result in a bias, which can be avoided by using hidden state models. All this provides a clear guideline for coding traits into characters. This paper gives practical examples of using the new framework for phylogenetic inference and comparative analysis.

---

Understanding the processes driving trait evolution is crucial for explaining evolutionary radiations (Price et al. 2010; Van Bocxlaer et al. 2010; Tobias et al. 2014), the origin of complexity and novelty (Moczek 2008; Ramirez and Michalik 2014), and for inferring phylogenies. For many of these analyses, we need to *(1)* discretize the trait (delimit the trait within a phenotype), *(2)* assess its primary homology (similarity), and finally *(3)* encode the trait (observations) into a character string or vector (see the definitions in Box 1, section A). This procedure, called *character construction* (Wiens 2001), is a basic stage of any analysis, and has a profound influence on all downstream stages. Despite the plethora of inference frameworks - be it parsimony, maximum likelihood or a Bayesian framework [reviewed in O’Meara (2012)] - the lack of repeatable and agreed-upon approaches for character construction generates considerable ambiguity. As a result, different hypotheses of discretization and different ways of coding the same hypothesis into a character may be proposed for the same trait (Hawkins et al. 1997; Strong and Lipscomb 1999; Ramirez 2007; Agnarsson and Coddington 2008). This disagreement naturally leads to inconsistent phylogenetic results [reviewed in Brazeau (2011)].

### Box 1. Definitions of the key terms.

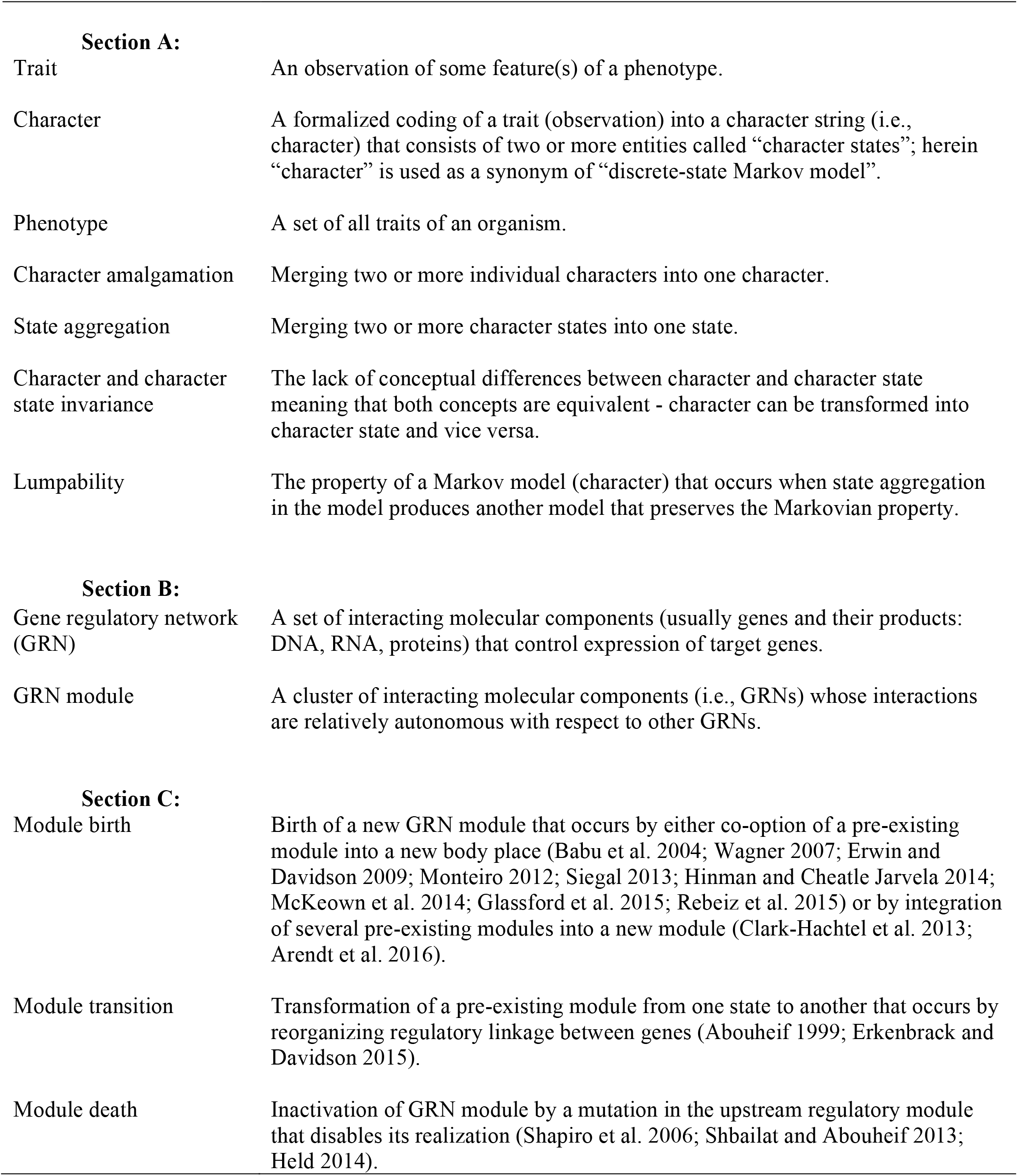

The ambiguity of character construction stems out from two key processes – hierarchical and hidden. The hierarchical process refers to the evolution of hierarchical relationships between traits that occur due to dependencies among anatomical body parts. For example, digits are located on limbs; loss of the limbs during evolution simultaneously triggers loss of the digits. The hidden process refers to the evolution of gene regulatory networks (GRNs) which underlay trait development (Wagner 2007; Carroll 2008; Houle et al. 2010); it implies that the actual driver of trait evolution is hidden from the direct observation of morphology. Evolution of developmental programs in organisms causes interactions between the hierarchical and hidden processes, making them into simultaneous drivers of trait changes. Unfortunately, the available phylogenetic methods do not accommodate these processes simultaneously. Thus, development of methods capable of concurrently modeling the two processes would automatically resolve much of the ambiguity associated with character construction. To tackle this problem, I propose a new integrative framework that uses the theory of structured Markov models [SMM, (Nodelman et al. 2002)], hidden Markov models [HMM, (Beaulieu et al. 2013)] and knowledge of organismal anatomies from anatomy ontologies. I assess the performance of the new framework using the two following case studies which fully characterize the problems of character construction.

1. *Hierarchical process: tail color problem and tail armor case*. – The ambiguity of coding anatomically dependent traits is best exemplified by the long-standing tail color problem (Maddison 1993; Hawkins et al. 1997) that seeks optimal scheme for scoring traits in species with no tails, blue tails and red tails. This problem has been widely discussed in parsimony literature but has not been solved (Maddison 1993; Hawkins et al. 1997; Strong and Lipscomb 1999; Brazeau 2011). I demonstrate that unlike parsimony, the proposed framework offers two natural solutions to this problem. To provide a better insight into modeling hierarchical dependencies, I also use a modified version of the tail color problem which I refer to as the “tail armor case”. This case considers tail trait that exhibits a two-level hierarchical dependency. Additionally, I stress the need for using anatomy ontologies to retrieve data about dependent traits and construct ontology-informed models.
2. *Hidden process: the two-scientist paradox*. – Discretization of phenotypic trait into states – the key step in character construction – commonly results in a mismatch between phenotypic and GRN state spaces. Since morphology is the product of GRNs, the failure to account for the hidden GRN evolution may bias phylogenetic analysis. I demonstrate this bias by reviewing the properties of GRN-to-phenotype maps in a Markov model context, and using a previously unknown issue called the “two-scientist paradox” – where the two scientists use different schemes to code the same trait and these schemes require different models to best fit the data due to the GRN-phenotype mismatch. I show that the bias associated with the hidden GRN evolution can be avoided by using the new framework and model selection procedure. From this modeling perspective, I discuss the confounding nature of the coding of a trait and the hidden factors underlying a trait’s evolution.

Even though the hierarchical and hidden processes may seem dissimilar, they are interacting due to the common mathematical machinery – modeling one requires use of the other. This interaction allows simultaneous modeling which, to a large extent, resolves all problems of character construction and provides a clear guideline for coding traits into characters. Additionally, present study gives practical examples of using the proposed framework for phylogenetic inference and comparative analysis.

This paper consists of the three main sections. The first section provides technical notes and theoretical background into SMM, HMM, and their key properties, which are essential for understanding the proposed models. The second section deals with the hierarchical process as illustrated by the tail problems, while the third section focuses on hidden process and the two-scientist paradox. The discussion, at the end, synthesizes recommendations for trait coding and modeling.

## 1. Theoretical background

This section includes five subsections dealing with the theory of Markov models. The first subsection begins with an overview of Markov models for trait evolution. The second one introduces the property of character and character state invariance that, in turn, is based on amalgamation and aggregation of Markov chains and their states (Box 1, section A). The third subsection overviews the techniques that can be used for modeling hierarchical dependencies. The forth subsection introduces the property of Markov model lumpability (Box 1, section A) that has a strong explanatory power for analyzing trait evolution. Finally, the fifth subsection deals with the ways to handle Markov models which violate lumpability property.

### 1.1. Overview of discrete-state Markov models for morphological data

#### Morphological character as a Markov model

The *traditional Markov model (MM)* implies that a character is a discrete-state and continuous-time Markov chain that moves sequentially from one state to another over the course of evolution. A Markov chain is defined by a transition rate matrix containing infinitesimal rates of change between the states, and an initial vector of probabilities at the root of a phylogenetic tree [for details see e.g., Huelsenbeck et al. (2003)]. This paper treats “Markov model” and “character” interchangeably, meaning that a rate matrix fully characterizes a character. The commonly used MM for a phylogenetic inference is the *Mk* model (Lewis 2001).

#### Structured Markov Models (SMM)

This class of models (Nodelman et al. 2002), also known as continuous-time Bayesian Networks (Shelton and Ciardo 2014), arises from traditional Markov models. The only difference between the two is that SMMs are equipped with a specific parameterization of the rate matrix to model dependencies. The application of SMM to phylogenetics was pioneered by Pagel (1994) to infer correlation between two binary characters. This paper extends the application of SMM for modeling various types of correlations and hierarchical relationships between anatomical parts, which have not been previously reviewed. In Supplementary Materials (Supplemental Materials are available on Dryad at https://doi.org/10.5061/dryad.j5r716b), I provide a separate set of *R* (Team 2008) functions that construct all types of SMMs reviewed in this paper and can be used to produce inputs for phylogenetic analyses.

#### Hidden Markov Models (HMM)

HMM elaborates traditional MM by splitting the model states into two layers - observable and hidden (Fig. 1a,c). The former represents the observable states of a phenotypic character, while the latter correspond to some unobserved factors influencing its evolution. The transitions between states are allowed only within the hidden layer; the observable states usually exhibit one-to-many mapping with the hidden states. If the mapping is one-to-one, then HMM collapses to MM. The use of HMM for the analysis of discrete traits has been recently pioneered by Beaulieu and O’Meara (2014) and Beaulieu et al. (2013).

**Figure 1.**
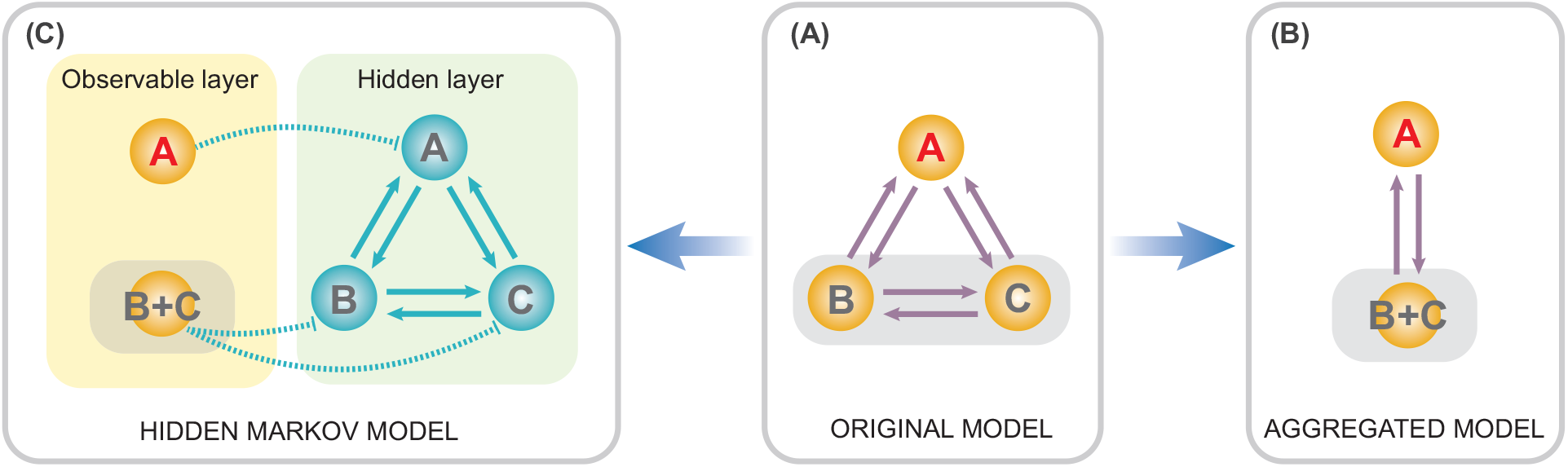
Hidden Markov model and lumpability. This original three-state Markov model (A) can be reduced to a two-state model by either directly aggregating the states (e.g., the states *B* and *C*) if the model is lumpable (B), or by using HMM with two observable and three hidden states if the model is not lumpable (C).

In the simulations used in this paper, I find it more flexible to construct HMM using ambiguous/polymorphic coding because, in model formalism, HMM and ambiguous/polymorphic coding are equivalent. For example, a HMM with three observable states {*absent, blue present (2), red present (3)*}, where *“absent”* includes two hidden states {*absent blue (0), absent red (1)*} is represented as a three-state character {*0&1, 2, 3*}; the first state is coded as polymorphic.

#### SMM+HMM

Equipping SMM with hidden states results in a more general class of structured Markov models that can simultaneously account for hierarchical and hidden processes. These models are the focus of the present paper.

#### Mk-SMM

The general SMM can be straightforwardly adopted for phylogenetic inference by converting it into an Mk-type model (Lewis 2001). This conversion is done by constraining its rate matrix to include one free parameter (that is usually interpreted as a branch length), setting its initial vector to contain equilibrium distribution at the root, and conditioning its likelihood on observing only variable characters in the data. The phylogenetic inference using various *Mk-SMM* is discussed below.

### 1.2. Invariance: character and character states are the same

A lack of general consensus on what constitutes a character versus character state presents a major challenge to the coding of traits. One group of studies insist that the distinction between the character and character state is important (Pinna 1991; Sereno 2007; Wagner 2015), while another suggests that both concepts are the same (Nelson and Platnick, 1981; Patterson, 1982). Here, I provide the evidence in favor of the latter statement.

#### Amalgamation of characters

The properties of SMM (Shelton and Ciardo 2014) allow mathematically valid amalgamation (Box 1, section A) of any number of characters thus removing distinction between character and character state. The amalgamation is especially straightforward when characters are independent (the dependent cases are reviewed in the next section). Suppose there are two characters: ***T*** – tail presence {with states: *absent (a), present (p)*}, and ***C*** - tail color {*red (r), blue (b)*}, which as we assume (for now) evolve independently according to the rate matrices:

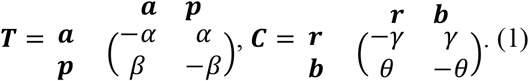

The rate matrix of the amalgamated character can be constructed via the following equation:

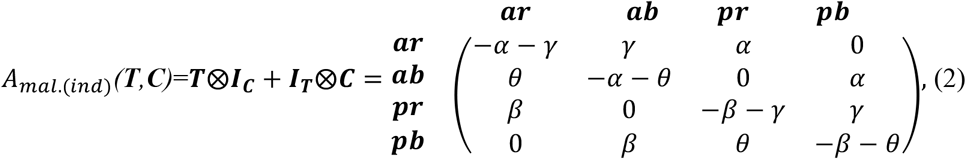

where ***I_T_*** and ***I_C_*** are the identity matrices for the two characters respectively, and ⊗ denotes the Kronecker product; hereafter I refer to this model as SMM-ind (for derivation see online Appendix S1). The four-state space of the amalgamated character is the state product (i.e. Cartesian product) of the initial characters that exhibits all combinations of ***T*** and ***C*** states, which are {*ar, ab, pr, pb*}. The zero elements in the matrix (2) prevent the initial states of ***T*** and ***C*** from changing simultaneously over infinitesimal interval of time since simultaneous change would imply a correlation between the characters (see the section Synchronous state change below). Also, these elements introduce a notion of state accessibility in phenotype (Stadler et al. 2001) that indicates which states are immediately accessible from present state through one-state change. To combine *n* independently evolving characters the equation (2) has to be successively repeated *n-1* times. For example, in the case of the three initial characters ***T, C***, and ***Z*** this means:

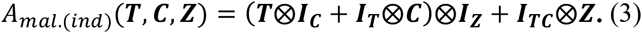

The matrices of amalgamated characters have peculiar symmetries - although matrix dimension grows fast, the vast majority of cells are zeros (Fig. 2c); the transition rates are located along the secondary diagonals. For a chain of *n* coevolving characters with the equal number of states *ω* the proportion of non-zero elements in the matrix is [1 + *n*(*ω* − 1)]/*ω^n^*. The number and pattern of secondary diagonals populated with transition rates increases with the number of states. If initial characters have equal number of states, the total number of secondary diagonals is *n*(*ω* − 1).

**Figure 2.**
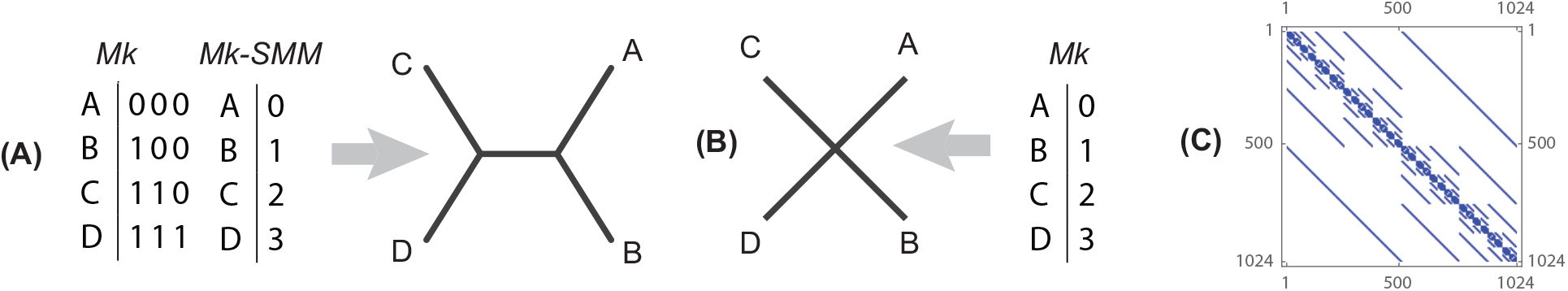
Character amalgamation. (A) Amalgamation of the three two-state characters evolving under *Mk* model results in one four-state character evolving under *Mk-SMM* (the amalgamation of the three two-state characters results in the eight-state rate matrix for *Mk-SMM;* in the given set of species only four states {0, 1, 2, 3} are observed, therefore the remaining four-states are omitted). The phylogenetic inference using these two datasets yields the identical and resolved topology. (B) If amalgamation of the three two-state characters into one four-state character is done without appropriately structuring the rate matrix (i.e. using four-state *Mk* model) then the topology is unresolved. (C) The amalgamated rate matrix (1024 states) of ten two-state characters; the zero cells (without rate values) of the matrix are shown in white.

#### Aggregation of states

The opposite of amalgamation is the aggregation of states, which allows decomposing one character into a set of several characters. The aggregation of states is mathematically valid only if Markov model is lumpable, which requires specific symmetries of rate matrix (see the Lumpability of Markov models) but always holds for independently evolving characters. The amalgamated character in the equation (2) can be decomposed with respect to the two partitions of its state space: 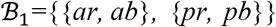 and 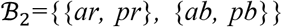, that gives transition matrices for the characters ***T*** and ***C*** respectively. This operation can be expressed as:

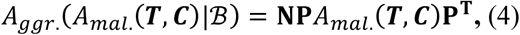

the meaning of the equation terms is provided in the section Lumpability of Markov models below.

To sum up, amalgamation combines separate characters into one single character, and aggregation allows the states of the same character to be represented as separate characters. Thus, regardless the initial way of discretizing organismal features into characters and states, the character and states are invariant (Box 1, section A) with respect to each other if rate matrices are appropriately structured. The simple simulation in Figure 2 exemplifies this theoretical consideration (online Appendix S2): the tree topology inferred using *Mk* model for the three two-state characters is the same as the topology when those three characters are amalgamated into one four-state character (Fig. 2a) and analyzed using structured *Mk* model (i.e., *Mk-SMM*) from the equation (3). If the four-state character is analyzed with traditional *Mk* model that is not properly structured, then the tree is unresolved since such amalgamation is invalid mathematically (Fig. 2b).

### 1.3. Modeling character dependencies using SMM

To the best of my knowledge, different coding schemes and trait dependencies generate three main types of correlation between characters: (i) a general type of correlation, (ii) “switch-on” dependency, and (iii) synchronous state change. The techniques of modeling these correlations for the two-character case are discussed below; and they can be extrapolated for an arbitrary number of characters using the equation (3).

#### General type of correlation

Independent evolution of characters ***T*** and ***C*** is defined by the rate matrix with the specific rate symmetries and the maximum of four free rate parameters [equation (2)]. Those symmetries can be relaxed by making all matrix rates different (Pagel 1994), which produces a complex pattern of correlation between the states (see online Appendix S3b). Such a matrix corresponds to the general type of correlation between the characters ***T*** and ***C*** (hereafter SMM-gen) that is:

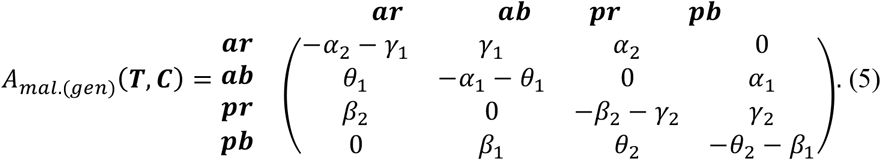

#### “Switch-on” type of dependency

This type arises due to anatomical dependencies between traits when hierarchically upstream trait switches on and off the downstream one. Consider two previous characters: ***T*** - tail presence {*a, p*}, and ***C*** - tail color {*r, b*}. Apparently, the tail color depends on the tail presence – both states of the character ***C*** are observable if and only if the character ***T*** is in the state present; if the tail is absent, then the color character is “switched-off’ and does not evolve. One of the ways to model such dependency is to amalgamate ***T*** and ***C*** as independently evolving using SMM-ind [equation (2)]. However, one may want to impose stricter relationships that reflect the anatomical hierarchy of the traits by prohibiting color changes when the tail is absent. It can be done using the modified version of the equation (2) that gives the following amalgamated rate matrix (hereafter SMM-sw):

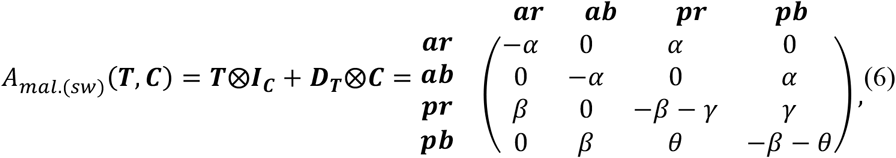

where ***D_T_*** is a diagonal matrix (online Appendix S3c). The difference between this matrix and that of SMM-ind is that here the transitions *ar → ab* and *ab → ar* are set to zero. In both SMM-ind and SMM-sw, the states *ar* and *ab* should not be necessarily interpreted as the “remainings of a pigment genetic machinery” that is capable to evolve even in the absence of the tail. Instead, they need be interpreted as the initial states for the tail color when the stochastic process switches from the *“tail absent”* to *“tail present”*.

#### Synchronous state change

This type frequently occurs when traits are redundantly coded using binary (absent/present) approach. Suppose there are two characters: (1) ***R*** – tail color red {*red absent (ar), red present (p_r_)*}, and (2) ***B*** - tail color blue {*blue absent (a_b_), blue present (p_b_)*}, defined by:

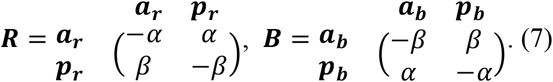

We assume that at a certain observation event, the tail can have only one color - either blue or red. This implies that if red is observed than blue is absent and vice versa. So, the states between ***R*** and ***B*** are mutually exclusive and hence change simultaneously over the course of evolution. In contrast to the previous amalgamation techniques, the synchronous evolution must allow only two-step transitions except for the transitions *p_r_p_b_ → a_r_a_b_* and *a_r_a_b_ →p_r_p_b_*, which are biologically impossible. This yields the following amalgamated rate matrix (hereafter SMM-syn):

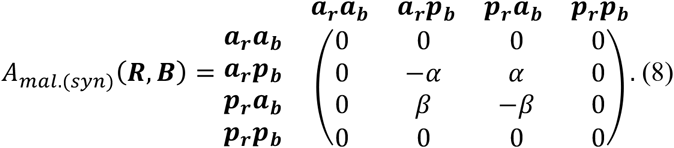

The two states *p_r_p_b_* and *a_r_a_b_* can be removed from this matrix as they are never visited by the Markov process. Their removal renders the matrix (8) to be absolutely equal to the two-state matrix in the equation (1) that defines the tail color.

### 1.4. Lumpability of Markov models

If aggregation of states in an original Markov model (character) produces an aggregated model that is still Markovian then the original model is called lumpable (Kemeny and Snell 1960; Rubino and Sericola 2014). The lumpability guarantees that transition rates and state sequence can be unbiasedly modeled using the aggregated model regardless the complexity of the original state space. If the aggregated model does not maintain the Markovian property, then the original model is not lumpable. The property of MM lumpability is essential for discretizing a trait into a character, maintaining character invariance, and modeling hierarchical and hidden processes. Below, I discuss conditions for the three types of lumpability which are relevant to trait modeling.

#### Strong lumpability

The aggregation of states is a partitioning of a rate matrix into partition blocks that correspond to the transition rates in the aggregated chain (Fig. 3). The strong lumpability implies that the original chain can be aggregated under any possible values of initial vector. The sufficient and necessary condition (hereafter, the row-wise sum rule, Figure 3) for a Markov chain to be strongly lumpable with respect to a given partitioning scheme is that the row-wise sum of rates within one partition block of rate matrix must be the same for all rows within the given partition block, and this property must hold for all blocks in the rate matrix (Kemeny and Snell 1960; Rubino and Sericola 2014). Suppose we want to lump an original rate matrix ***M*** using a partitioning scheme 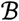 into the aggregated matrix 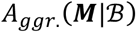. If the model is lumpable the row-wise sum rule implies the following equality to hold (Kemeny and Snell 1960):

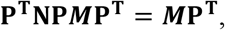

where **N** and **P** are the matrices specifying state assignment between the initial and aggregated models (online Appendix S3a). So, the rates in the aggregated matrix 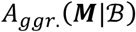 can be expressed as:

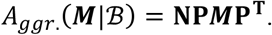

**Figure 3.**
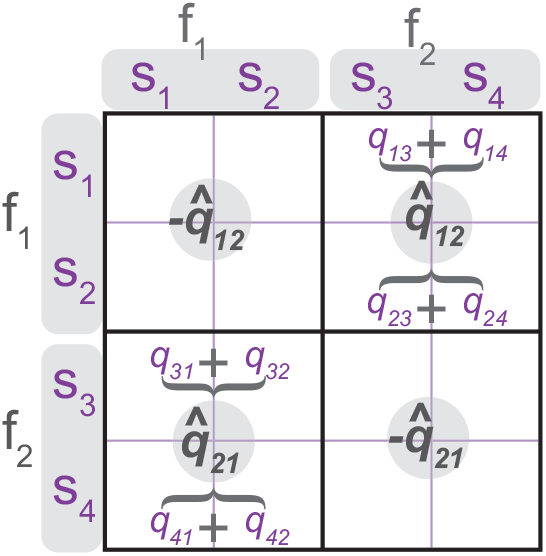
Lumpable Markov model. Symmetries in the rate matrix that are necessary for strong lumpability (row-wise sum rule); the original four-state {*s_1_, s_2_, s_3_, s_4_*} matrix with rates *q_ij_* is aggregated into the two-state {*f_1_* = { *s_1_, s_2_*}, *f_2_* = {*s_3_, s_4_*}} matrix with rates 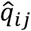; the aggregation is lumpable iff 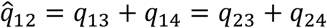 and 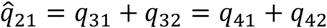.

For example, consider the four-state rate matrix *A_mal.(ind)_(**T,C**)* from the equation (2). Its states can be aggregated using the partitioning schemes 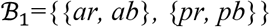 that results in the character ***T*** from the equation (1). The aggregation is possible because the row-wise sum rule is satisfied that implies the following equalities:

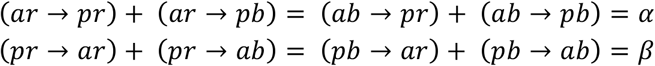

Here, the rates *α* and *β* on RHS define precisely the rates of the aggregated character ***T***. The partitioning scheme 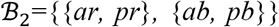 also produces a lumpable model resulting in the character ***C*** from the equation (1). In contrast, SMM-gen from the equation (5) cannot be lumped since its rate matrix, with all rates different, violates the row-wise sum rule under any partitioning scheme.

#### Weak lumpability

Weak lumpability allows lumping the original chain only under particular values of the initial vector (Kemeny and Snell 1960; Rubino and Sericola 2014), which imposes stricter dependencies between the initial vector and rate matrix. Generally, these dependencies cannot be found analytically but a finite algorithm can be used to elucidate them (Rubino and Sericola 1993, 2014). An interesting case of weak lumpability arises when the initial vector of a Markov chain contains an equilibrium distribution, meaning that the probability of observing chain’s states after some time has the same equilibrium distribution. Such chain is weakly lumpable with respect to any possible partitioning scheme (online Appendix S4). Although Markov models that have the initial vector with equilibrium distribution are commonly used in phylogenetics, the weak lumpability should not be expected to hold due to the way Felsenstein’s pruning algorithm (Felsenstein 1981) calculates likelihood (online Appendix S4). Thus, weak lumpability is omitted from the further discussion in this paper.

#### Nearly lumpable chains

This type of lumpability can occur in a large multistate character, thereby allowing to lump it with insignificant error. Such a character can be constructed by amalgamating many elementary characters into one large rate matrix using SMM-gen (see General type of correlation). Aggregation of states in such matrix can be thought of as mapping evolutionary processes occurring at the level of DNA sites or numerous GRNs to its realization at the phenotypic level. For example, consider a DNA locus of 1000 sites with each site being a four-state character (four nucleotides). All sites can be amalgamated into one large character with 4^1000^ states; this character possess peculiar symmetries of the rate matrix(e.g., see Fig. 2c) in which over 99% of cells are set to zero (see the section Amalgamation of characters). Suppose that those 4^1000^ states of the DNA character are mapped only to a few states of a phenotypic character. Since the original state space is significantly larger, each state of the phenotypic character is an aggregation of many original DNA states.

In a trivial case of equal evolutionary rates across all sites, the strong lumpability condition can be satisfied for numerous partitioning schemes that can be applied to amalgamate the DNA character. In contrast, if the amalgamation is constructed using SMM-gen with all rates different to reflect the most complex scenario of correlated evolution, then, obviously, such character drastically violates the condition of strong lumpability. However, if rates in the rate matrix, and values of the initial vector are identically and independently distributed (*i.i.d.*), which does not rule out the possibility they are different, then the amalgamated DNA character is nearly lumpable under any possible partitioning scheme. This occurs because the *i.i.d.* condition produces an aggregated chain whose error, in approximating the original rates, is insignificant as the number of the original states increases, while the number of the aggregated states decreases (online Appendix S5).

### 1.5. Handling non-lumpable models

If the aforementioned conditions of lumpability are violated, then the aggregated model is non-Markovian. Thus, the inference of the aggregated process using a traditional MM is biased. Hidden Markov models are a flexible tool to overcome non-lumpable aggregation by treating the original and aggregated processes as hidden and observable layers of the same model, respectively (Fig. 1). The application of this technique is considered in the following example.

#### Modeling SMM using HMM

The amalgamation of the tail characters via SMM-ind and SMM-sw [see the equations (2, 6)] produces matrices with four states {*ar, ab, pr, pb*}. The states *ar* and *ab* correspond to the same observation specifying the absence of the tail since tail color cannot be observed when the tail is absent. Notably, these matrices, as might be expected, cannot be reduced to a three-state matrix with states {*a, r, b*} under any values of their rate parameters because they are not lumpable under the partitioning scheme {{*ar, ab*}, {*pr*}, {*pb*}} (online Appendix S6). Thus, modeling SMM-ind and SMM-sw requires using HMM that includes three observable states {*a, r, b*}, and the observable state *a* consists of the two hidden states *ar* and *ab*.

## 2. Modeling hierarchical process: anatomy ontologies and structured Markov models with hidden states

This section discusses the principles of modeling morphologically dependent traits. It starts with the overview of the tail color problem and tail armor case. Next, it demonstrates how SMM + HMM can be used to solve these issues and assesses the SMM + HMM performance using simulations. Finally, it emphasizes the need of using ontology-informed models for modeling dependent traits.

### 2.1. Review of the tail color problem and tail armor case

Even if discretization is straightforward, anatomical dependencies render trait encoding into character ambiguous due to the missing consensus about which coding approach to use (Maddison 1993; Pleijel 1995; Hawkins et al. 1997; Strong and Lipscomb 1999; Forey and Kitching 2000; Brazeau 2011). The tail color problem (Maddison 1993) exhibits a typical case of coding an anatomically-dependent trait in species with no tail, a blue tail and a red tail; thus, a solution to it could be extrapolated to all other cases with dependencies. Four main schemes have been traditionally proposed to encode the tail traits (Fig. 4). These schemes differ in the number of characters and states that characterize the dependencies. The *first scheme* (Fig. 4a) employs two characters: (i) tail presence, and (ii) tail color; the absence of the tail in the second character is coded as the inapplicable observation. (i.e., when inapplicable character state is coded with “–” which is interpreted as a missing entity by available methods). The *second scheme* (Fig. 4b) is similar to the previous one but encodes tail absence as a separate state. The *third scheme* (Fig. 4c) employs three binary characters: (i) tail present, (ii) blue tail present, and (iii) red tail present. Finally, the *fourth scheme* (Fig. 4d) uses one three-state character.

**Figure 4.**
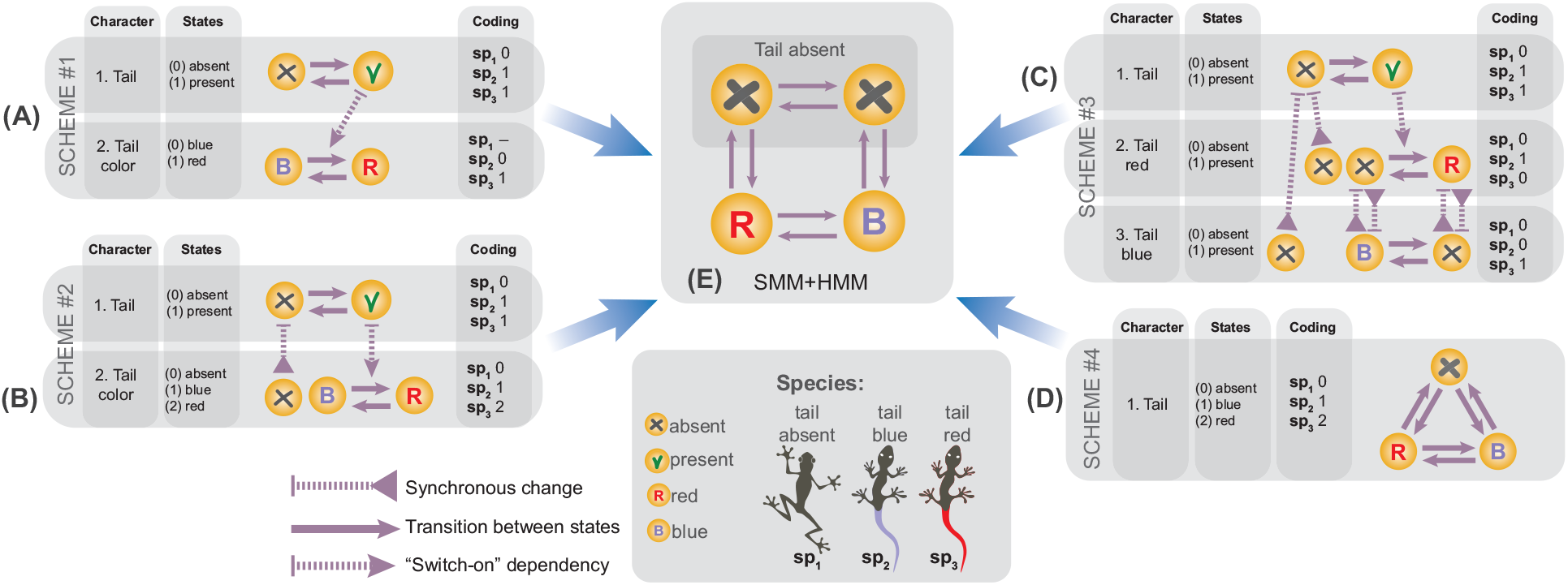
The coding schemes for the tail color problem. (A-D) The four alternative schemes for coding the tail traits; a graph next to each scheme shows dependencies between the characters, which arise due to the anatomical dependencies between the traits. If each of the four schemes is amalgamated into one character with the appropriate incorporation of those dependencies, then all schemes collapse to the same structured Markov model (E).

The behavior of these schemes have been intensively assessed over years in the context of parsimony but all of the schemes have been shown to have shortcomings (Maddison 1993; Hawkins et al. 1997; Strong and Lipscomb 1999; Brazeau 2011). Specifically, the schemes #2 and #3 fail to provide biologically logical character optimization, and the scheme #4 does not include hierarchical information and thereby fails to reconstruct complex dependencies.

Currently, the scheme #1 (inapplicable coding) is accepted to be the most optimal solution to the problem; however, it is known to suffer from undesirable behavior. Maddison (1993) gave the following example to demonstrate it. Suppose there is a tree of 14 species (Fig. 5a) where the tailed species are nested within the left and right clades of the tailless species. The tree is assumed to be fully resolved except for the relationships of the left tailed clade (LTC in Fig. 5b). Next, Maddison shows that one of the parsimonious resolutions of the LTC is identical to that of the right tailed clade (Fig. 5c). However, if the red and blue tailed species in this clade are swapped, then the resulting tree is no longer parsimonious (Fig. 5d). This means that individual tailed clades influence each others’ parsimony score, even though they are widely separated on phylogeny. Such effect is inappropriate since the evolution of tail color in these clades should be considered in isolation, and hence these two resolutions (Fig. 5c,d) must have equal parsimony scores.

**Figure 5.**
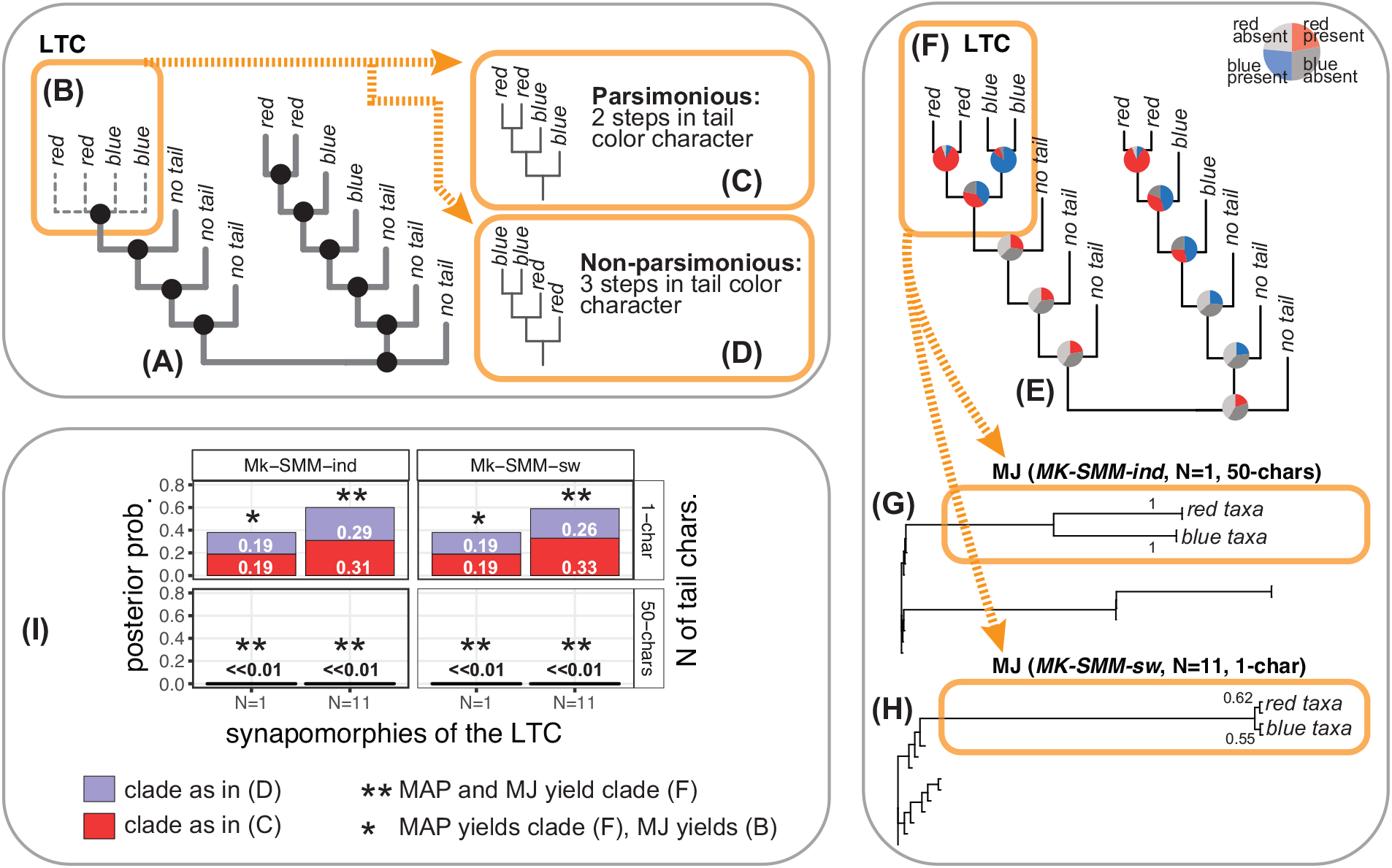
The tail color problem. (A) The tree from Maddison (1993) that exemplifies the tail color problem where the tailed species are nested within the two major clades of the tailless species; all relationships are assumed to be resolved (black balls) except those in the left tailed clade (LTC) shown in (B). (C, D) Two possible resolutions of the clade (B) when the coding scheme #1 is used: (C) – parsimonious resolution; (D) – non-parsimonious resolution. (E) MAP (maximum a posteriori) tree with the reconstructed ancestral states from the analysis using *Mk-SMM-ind*, one tail character (1-char), and only one synapomorphy (N=1) supporting the LTC. (F) One possible resolution of the LTC. (G, H) MJ (50% majority rule) trees from *Mk-SMM-ind* and *Mk-SMM-sw* analyses with the LTC resolved as shown in (F); numbers indicate posterior probabilities of the subclades in (F). (I) Plot of the posterior probabilities of the clades shown in (C) and (D) across all eight analyses. *Abbreviations: N* – number of synapomorphies (1 or 11) supporting the LTC; *x-chars* – number of tail characters (1 or 50).

Below, I demonstrate that SMM offers two natural solutions to the tail color problem which do not suffer from any of the above-mentioned shortcomings. Also, I use the modified version of the tail color problem – the “tail armor case” (Fig. 6d-e) – to demonstrate that SMM can be flexibly used to model any complex hierarchical relationships. The tail armor case considers four species that possess a tail with blue or red armor, a tail without armor, and no tail; thus, it implies a two-level dependency – armor color depends on armor presence that, in turn, depends on tail presence (Fig. 6e). Finally, I theoretically assess the two solutions in the light of the alternative coding schemes and ontology-informed models.

**Figure 6.**
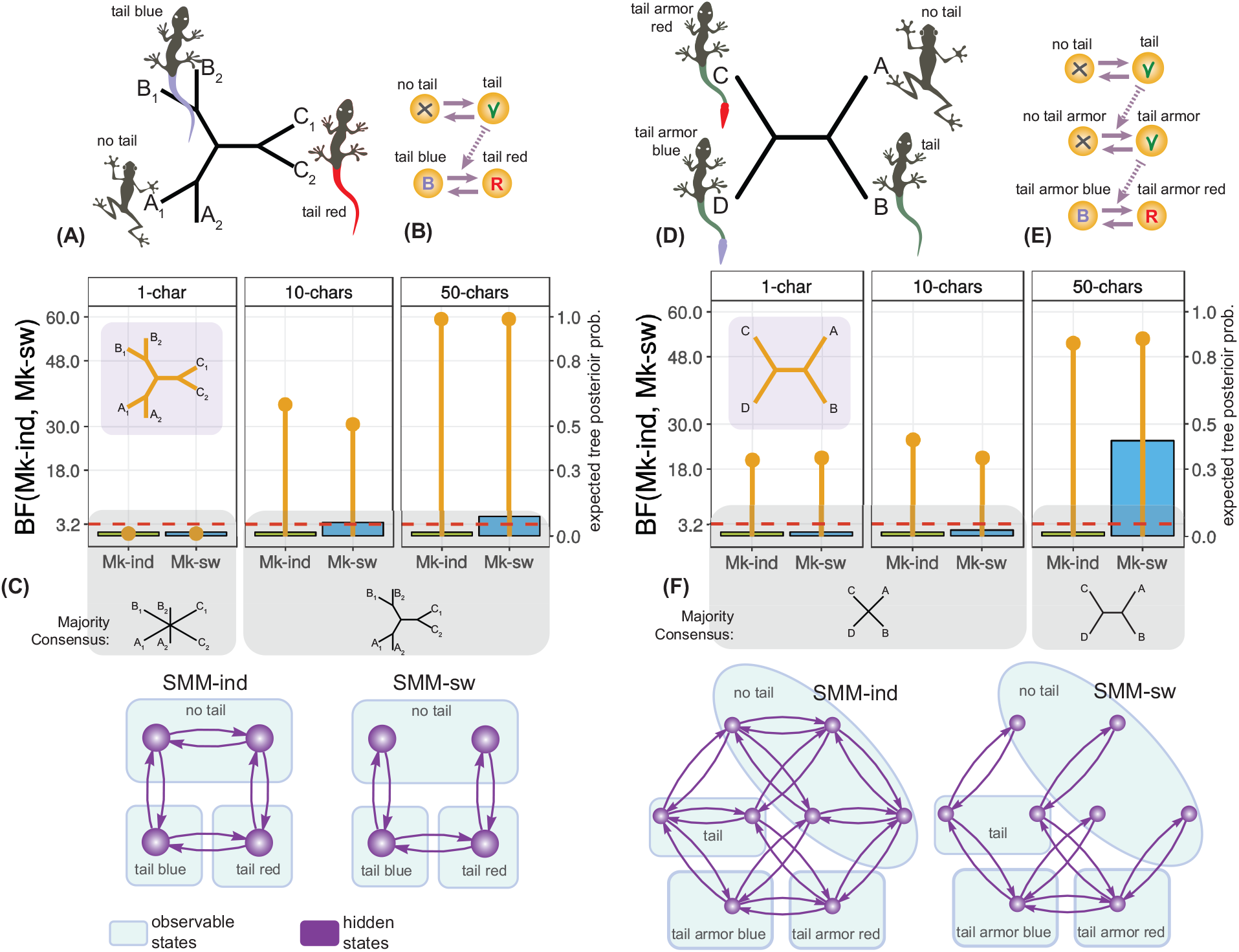
Model comparison for the tail problems. (A-C) Tail color problem. (D-F) Tail armor case. (A,D) Expected trees. (B,E) Graphs depicting anatomical dependencies between the traits. (C,F) Results of inference; *Mk-ind* and *Mk-sw* correspond to one-parameter *Mk-SMM-ind, Mk-SMM-sw* respectively; for each model the column bar indicates BF value (left-side legend), while thin bar indicates the posterior probability of the expected tree (right-side legend); note that the higher BF values indicate worse model fit, the threshold of BF<3.2 means that the two models yield similar fit. The graphs on the bottom show the topologies of SMM models used for the inference.

### 2.2. Solution to the tail color problem and tail armor case

In the context of Markov models, the tail traits and their dependencies can be naturally modeled with the two main approaches [see the equations (2, 6)]: (i) amalgamating the two characters as independently evolving via SMM-ind, or (ii) amalgamating the two characters through the “switch-on” dependency via SMM-sw. Both SMM-ind and SMM-sw represent the solutions to the tail color problem, tail armor case and any general case of anatomical dependency.

The phylogenetic inference with SMM-ind and SMM-sw can be performed by converting them into *Mk*-type models with one-parameter (i.e., branch-length): *Mk-SMM-ind* and *Mk-SMM-sw* respectively (see Overview of discrete-state Markov models for morphological data, and online Appendix S7). Despite having identical observable state space, *Mk-SMM-ind* and *Mk-SMM-sw* imply different assumptions for trait evolution. In analyses, these models can be compared using statistical methods for model selection (e.g., Akaike information criterion, Bayes factor, etc.). The use of model selection methods becomes possible due to the identity of their state spaces that keep the data the same.

### 2.3. Demonstrative simulations

I assess the performance of *Mk-SMM-ind* and *Mk-SMM-sw* in the Bayesian framework using *RevBayes* (Höhna et al. 2016) by running the two series of simulations (see the scripts in Supplementary Materials). The first series tests the model behavior using the original formulation of the tail color problem (hereafter, *tail color problem (TCP) simulations*); the second series tests the ability of the models to account for complex hierarchical dependencies (hereafter, *hierarchical dependencies (HD) simulations*). In all simulations, the tail color character was coded with three observable and four hidden states states, while the the tail armor character was coded using four observable and eight hidden states (online Appendix S7-8, Figure 6).

#### 1a. TCP simulations: methods

The original conditions given in Maddison (1993) for the tail color problem are followed to show that SMMs are not subjected to the inappropriate statistical behavior of the other coding schemes. All relationships of the fourteen-species tree (except those within the LTC) were constrained to be resolved, and supported by, at least, one binary character to avoid zero-length branches. In this set-up, *RevBayes* samples possible topologies of the LTC from the posterior distribution; I use this sample to assess the posterior probability of the alternative resolutions. For each model – *Mk-SMM-ind* and *Mk-SMM-sw* – I run four simulations under the combinations of the following conditions: (1) varying number of tail characters (one or fifty) to assess the effect of information content on model behavior, and (2) varying number of synapomorphies supporting the LTC (one or eleven characters) to assess the effect of branch length on the topology.

#### 1b. TCP simulations: results and discussion

All simulations for *Mk-SMM-ind* and *Mk-SMM-sw* show that posterior probabilities for the two alternative resolutions of the LTC (Fig. 5c,d) are (almost) identical (maximum difference is 0.05, see Figure 5i). This clearly indicates that the behavior of SMM is drastically different from that of parsimony algorithms with inapplicable coding. In contrast to parsimony that favors the clade in Figure 5c over the clade in Figure 5d, SMM equally samples the two alternative resolutions from the posterior distribution. Thus, in SMM the resolution of the right tail clade does not notably affect that of the LTC. Yet another notable aspect – the strict consensus of the parsimony analysis produces the undesirable topology for the LTC (as the one showed in Figure 5c). In contrast, SMM tends to resolve the LTC by supporting the monophyly of red and blue tailed subclades respectively (Fig. 5f); this resolution was recovered in all analyses on maximum a posteriori trees (Fig. 5i,f), and in all analyses on majority rule (50%) consensus (MJ) trees (Fig. 5i,g-h) when fifty tail characters or eleven synapomorphies were used. The analyses where the LTC was supported by only one tail character (without any extra synapomorphies) yielded unresolved LTC as in Figure 5b. This occurs because the performance of SMM, unlike parsimony, also depends on branch length. If information content in the branch length is sufficient (e.g., when the eleven synapomorphies are used to support the LTC), then MJ can fully resolve the LTC even in the presence of only one tail character (Fig. 5h).

#### 2a. HD simulations: methods

These simulations evaluate *Mk-SMM-ind* and *Mk-SMM-sw* in the context of model selection and their ability to model hierarchical dependencies. The inference was performed for two datasets (the modified tail color and tail armor cases) using one, ten, and fifty identical characters. The different number of characters was chosen to evaluate the effect of data information on model behavior. The modified dataset for the tail color problem included only three pairs of species - each pair had the same character pattern (Fig. 6a-b); these six species were chosen to assess topologies of unrooted trees (there is only one unrooted tree for three taxa). The dataset for the tail armor case included four species which was sufficient to assess the two-level hierarchal dependency (Fig. 6d-e). The relative model performance was evaluated using Bayes factors (BF) which compares the ratio of marginal likelihoods of two candidate models (Lavine and Schervish 1999). To interpret BF threshold for which one model shows better fit than the other, I use the scale proposed by Jeffreys (1961). In this scale the BF value <3.2 indicates equal fit for two models; BF>10 suggest strong support in favor of the first model. The marginal likelihood was calculated using stepping-stone (Xie et al. 2011) and path-sampling approaches (Lartillot and Philippe 2006) implemented in *RevBayes*. Since both methods yielded similar values (i.e., BF<3.2), only the former is used in the discussion below.

#### 2b. HD simulations: results and discussion

The expected topology for the modified tail color dataset must group together the pairs of species with the identical characters (Fig. 6a). Both *Mk-SMM-ind* and *Mk-SMM-sw* yield the majority consensus tree to be the same with the expected topology when datasets include more than one character (Fig. 6c). The inference with one character does not differentiate between the candidate models (all BF values <3.2); however, as the character number increases to fifty, *Mk-SMM-ind* yields a moderately better fit than *Mk-SMM-sw* [BF(*Mk-SMM-ind, Mk-SMM-sw*) = 5.4]. In the armor color case, the intuitive tree is expected to group the species with armor in one clade, and those without armor in another--thus reflecting the putative evolutionary sequence of tail and armor emergence (Fig. 6d). All analyses with one and ten characters yield unresolved majority consensus, and similar fit for the models (Figure 6f). In the analyses with fifty characters, the two models produce the expected consensus but *Mk-SMM-ind* reveals a significantly better fit [BF(*Mk-SMM-ind, Mk-SMM-sw*) = 25.6].

#### Simulations: general discussion

To summarize, the simulations above demonstrate that dependent traits can be efficiently modeled by SMMs with hidden states. These models –*Mk-SMM-ind*, and *Mk-SMM-sw* – do not display the inappropriate behavior detected by parsimony, and can be differentiated using model selection criteria. The better fit of *Mk-SMM-ind*, in both sets of the simulations, likely occurs due to the simplicity of the example datasets (*Mk-SMM-sw* describes more complex and constrained relationships). Both models reveal the expected tree when the amount of data is sufficient; however, the topological performance of the two models should not be expected to be the same in complex datasets. Thus, similar to DNA data, model selection for morphology-based inference is important.

### 2.4. Comparison of the coding schemes against SMM-ind and SMM-sw

Let us now assess the four alternative coding schemes (Fig. 4) against the presented solutions that are based on SMM-ind and SMM-sw. To make the coding schemes comparable, the variable number of characters between them has to be eliminated by using the character invariance property and amalgamating the characters of each scheme into one single character (online Appendix S9).

#### Scheme #1

Amongst all alternative schemes only the scheme #1 {amalgamated states: *ar, ab, pr, pb* }--which uses inapplicable coding--is biologically meaningful. It exhibits exactly SMM-ind that models the tail traits as independent initial characters. This solution has been known and widely applied in phylogenetics before. As shown above, in parsimony this solution suffers from the undesirable effect which avoided by using SMM. The rate matrix for the scheme #1 can be further elaborated using the “switch-on” dependency to yield the second solution via SMM-sw that has not been previously reported.

Note, the SMM state space cannot be reduced to the three states {*a, r, b*} [see Handling non-lumpable models]. However, SMM-ind is lumpable with respect to the tail and color characters [see the equation (4)] that precisely matches the scheme #1. This does not hold for SMM-sw whose rate matrix cannot be lumped in the same manner (online Appendix S6); thus, SMM-sw can be modeled only using the hidden states.

#### Scheme #4

This scheme (single three-state character), having at first glance similar state space, is quite different from the scheme #1. For comparison, let us expand the state space of the scheme #4 to make it identical to that of the scheme #1. This implies creating a four-state rate matrix that, if lumped, collapses to the original three-state one (online Appendix S10a). In contrast to the scheme #1, the expanded matrix of the scheme #4 has non-zero rates for two-step transitions, thus failing to account for the hierarchical dependencies and hence lacking the notion of state accessibility in phenotype (Stadler et al. 2001). If this scheme is used to model complex dependencies (e.g., as those in the tail armor case), then the resulting topology would be completely unresolved (e.g., the topology for “1-char” in Fig. 6f) which was also observed in parsimony (Hawkins et al. 1997).

#### Schemes #2&#3

The amalgamated state space of the schemes #2 and #3 differs significantly from the rest by having spurious states. For example, the state of scheme #2 *“tail present, no color”* and the state of scheme # 3 *“tail present, tail blue, tail red”* are logically contradictory since the tail is colored once it appears, and tail cannot be blue and red at the same time. During phylogenetic inference, these states will be reconstructed as ancestral conditions which will make the inference biologically misleading as was previously noted for parsimony (Forey and Kitching 2000). Even though, the use of the schemes #2 and #3 may result in a “correct” topology, these schemes should not be used for coding as their biological interpretation is misleading.

#### Character coding invariance

As shown above, SMM-ind and SMM-sw represent two solutions for the tail color problem, which can be coded with the scheme #1. The schemes #2, #3 and #4 fail to meaningfully accommodate hierarchical relationships because they contain synchronous changes, redundant states or improperly structured rate matrices (see the dependency graphs in Figure 4) that cannot be modeled directly through SMM-ind or SMM-sw. In fact, the amalgamation of the schemes #2, #3 and #4 has to be derived differently by considering their specific dependencies and using SMM-syn. Interestingly, this derivation makes the schemes #2, #3 and #4 identical to the scheme #1 (online Appendix S10). Thus, the alternative coding approaches become invariant (i.e., equivalent) with respect to each other if anatomical dependencies between traits are appropriately incorporated using SMM.

### 2.5. Ancestral state reconstruction on a known tree

The provided considerations for hierarchically dependent traits hold for both phylogenetic inference and ancestral character state reconstruction. Using inapplicable coding (i.e., the scheme #1), that exhibits one of the appropriate ways for coding hierarchical trait, has been common in tree inference. This is in contrast to ancestral character state reconstruction on a known tree where the use of one character with multiple states would be a common choice (i.e., the scheme #4). As was shown earlier, this is a misleading approach since SMM-ind and SMM-sw cannot be substituted by a three-state rate matrix. Thus, I suggest using hidden state models (i.e., SMM-ind and SMM-sw) for ancestral reconstruction whenever anatomical dependence in traits are detected. In ancestral reconstruction, the rate constraints in SMM-ind and SMM-sw can be relaxed by, for example, making all rates different as those in SMM-gen.

### 2.6. Ontology-informed structured Markov models

The hierarchical dependencies between traits are imposed by the structure of organismal anatomy and hence can be retrieved using anatomy ontologies. This is turning ontologies into a promising tool for arranging, querying and managing anatomical data (Deans et al. 2015). The need for integrating ontologies with morphology-based phylogenetic analysis has been recently emphasized and discussed (Vogt 2017a, 2017b, 2018). Unfortunately, so far, anatomy ontologies are available only for several taxa [e.g., Mungall et al. (2012) and Yoder et al. (2010)]. Nevertheless, the available software allows direct linking of character matrices with anatomy ontologies (Balhoff et al. 2010). This provides an opportunity for an automatic extraction of anatomical dependencies from ontologies linked to characters (Dececchi et al. 2016). Alternatively, dependencies can be incorporated by simply using a scientist’s own knowledge of organismal anatomy. Integration of SMM with ontologies opens possibilities for reconstructing ancestral anatomy ontologies in a way similar to the parsimony-based method proposed by Ramirez and Michalik (2014). In this light, a character becomes not merely a vector of numbers (i.e., states) but a graph reflecting various relationships, each with its own anatomical meaning. Such graph, produced by linking the tail character with UBERON ontology (Mungall et al. 2012), is shown in Figure 7. Structured Markov models can be used to infer topology of such graphs as well. It can be anticipated that further developments in this field will lead to a creation of an automated pipeline for constructing ontology-informed SMM+HMM models. This pipeline will open up new perspectives for modeling evolution of entire organismal anatomies.

**Figure 7.**
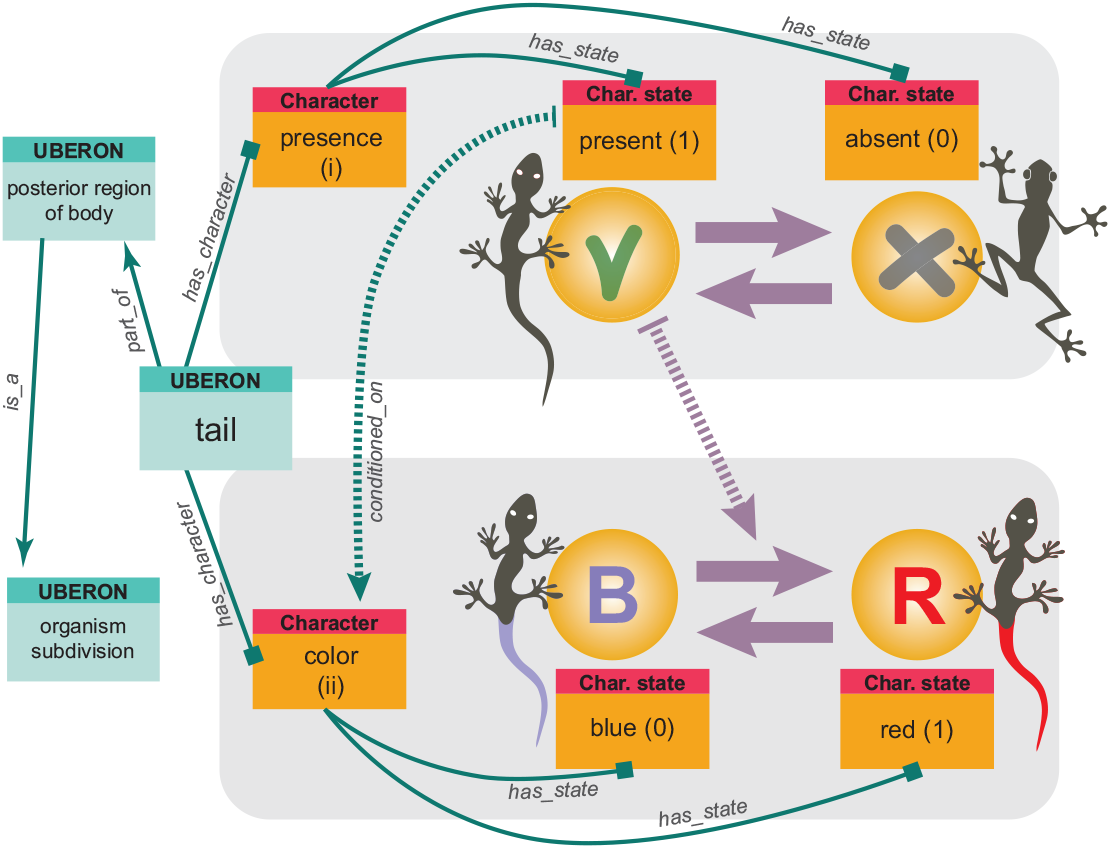
Ontology-informed character. The tail characters linked with ontology. The links (i.e., arrows) show various types of ontological relationships between the characters and between entities of UBERON anatomy ontology.

## 3. Modeling hidden processes: morphology, GRNs and structured Markov models with hidden states

Besides the hierarchical process reviewed in the previous section, the hidden process is another key driver of trait evolution. Phenotypic traits are the products of realization of GRN modules (see Box 1 section B) over spatiotemporal scales of embryo development. Thus, GRN modules can be reasonably considered to be the elementary hidden units in the genotype-to-phenotype map (Pigliucci 2010). In this respect, it is essential to summarize general mechanisms of GRN evolution, their affects on traits, and properties of GRN-to-phenotype (GRP) maps. Therefore, this section starts with an overview of the properties of GRP maps which are critical for understanding their modeling principles. Next, it shows that in most cases GRP maps cannot be unbiasedly modeled using MM due to the violation of lumpability property; to overcome this problem HMM must be used. Finally, to demonstrate the use of HMM, this section assesses the hidden process from the perspective of the two-scientist paradox.

### 3.1. Overview of correspondence between GRNs and morphology

To assess the properties GRP maps let us formalize the evolution of GRNs using the framework of Markov models. In this regard, each step in the evolutionary path of a GRN module is a state of a Markov chain. The complete evolutionary path of the module involves: module birth, the transition between module states, and module death (Box 1, section C). GRP map can be viewed as a mapping of the states of GRN Markov model to those of a discrete character. This mapping falls into three main types overviewed below (Fig. 8).

**Figure 8.**
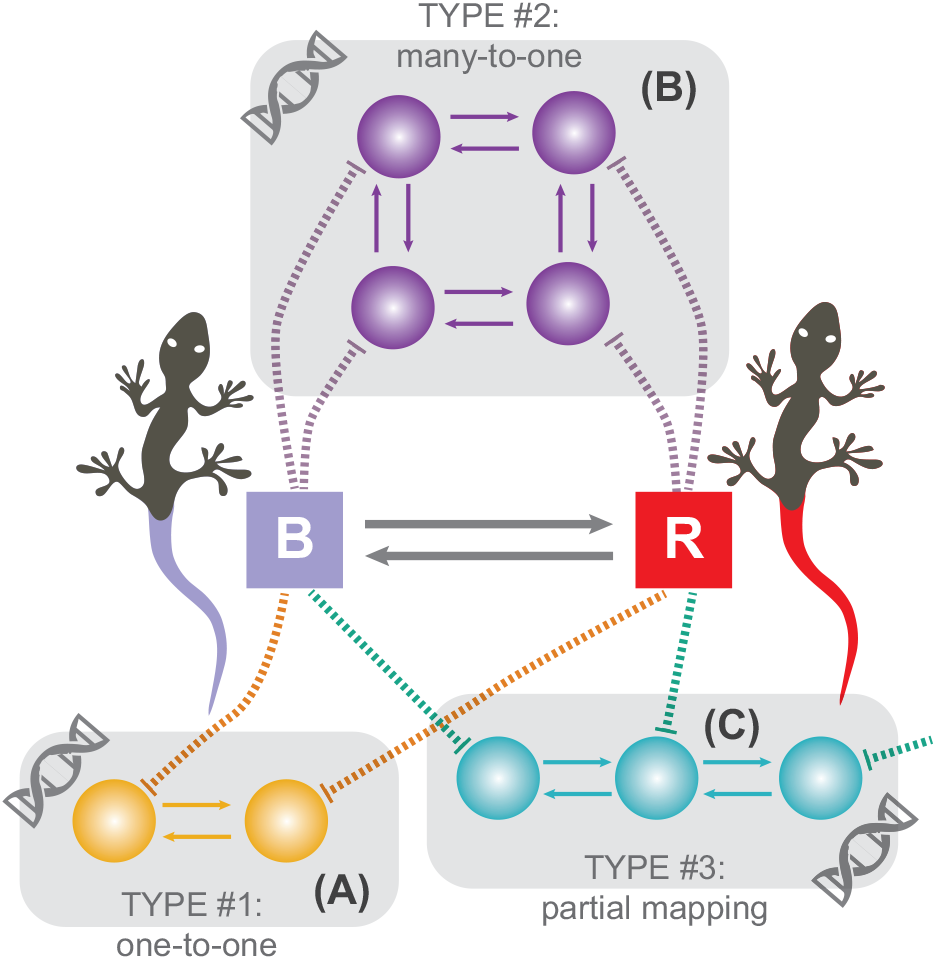
GRN-to-phenotype mapping. The tail color trait comprises two states: blue and red (shown with rectangles “B” and “R” respectively). The states of hypothetical GRN modules that produce the tail color trait are shown with spheres. (A) One-to-one mapping. (B) Many-to-one mapping. (C) Partial mapping.

#### Type 1: one-to-one

One-to-one mapping between states of GRN modules and those of a discrete character indicates that primary homology hypothesis correctly identifies underlying genetic space (Fig. 8a). This case is ideal but rarely realistic since morphological traits are usually the products of complex gene interactions.

#### Type 2: many-to-one

Many-to-one mapping is a scenario when several GRN states are mapped onto one trait state (Fig. 8b) indicating that the trait is controlled by multiple genetic factors which are usually unknown to a researcher (Rebeiz et al. 2015). Numerous evo-devo studies show that this scenario is a commonplace (Abouheif 1999; Hall 2003; Moczek 2008; McCune and Schimenti 2012; Wagner 2015). For instance, some males of *Drosophila* have a pigmented spot, located on the wing, that they use in a courtship display (Prud’Homme et al. 2006). This spot is controlled by one gene, *yellow*, and has similar shapes across different species. For a phylogenetic analysis, it would be reasonable to code this trait using a two-state character {*spot present, spot absent*}, which was done in the study of Prud’Homme et al. (2006). That study revealed multiple gains of the spot during the evolution of *Drosophila*. Interestingly, these independent gains were caused by a different mechanism – the co-option of different *cis*-regulatory elements associated with *yellow*. Evidently, even a seemingly simple trait such as the wing spot may, in fact, have a complex GRN state space.

Another situation when many-to-one correspondence can be a commonplace refers to the incompleteness of morphological examination that may arise when external structures are examined without reference to skeletal structures (in vertebrates), or when external skeletal structures are examined without reference to underlying muscles (in invertebrates). For example, different lineages of salamanders have similarly elongated body shapes which may be hypothesized to be homologous assuming that the phylogeny is unknown. However, the mechanism of body elongation varies across the lineages by adding and extending different individual vertebrae (Wake et al. 2011). If body shape is studied without reference to the skeleton, then similarly elongated bodies would be scored with the same character state. However, this approach is misleading since the different mechanism of elongation may produce the same shapes, which reflects many-to-one mapping between the underlying state space and observation. Thus, in the model formalism, this and the *Drosophila* case are the same.

#### Type 3: partial mapping

The partial mapping occurs when one trait is controlled by interdependent GRN modules but this interdependence in not reflected in the coding--meaning that the GRN space producing the trait is only partially identified (Fig. 8c). For example, the aforementioned tail color problem can be roughly formulated in the GRN terms using the two modules: one controlling tail development, and another, nested module, controlling color; considering the color character as a separate entity (i.e., using the coding schemes #2-#4) results in this type of mapping because tail color and tail presence are the parts of the same developmental process; thus, they have to be treated as one character. The partial mapping may be a consequence of pleiotropy and epistasis when seemingly unrelated traits are correlated due to the shared genetic factors. Moreover, in the model formalism, partial mapping corresponds to any type of correlation that is missed due to a spurious trait discretization or model choice. For example, traits of insect mouthparts can undergo simultaneous evolutionary changes when species get adapted to new feeding conditions. In this situation, treating different elements (e.g., mandibles, labrum, etc.) of the mouthparts as separate characters is misleading; the dependencies have to be incorporated into the model to reflect the correlated evolution. The diversity of various types of partial mapping occurs because, in phylogenetics, they all match the same general model – SMM-gen. In some cases, the existence of correlation is obvious and can be retrieved from anatomy ontology; however, cases with unobservable correlation - when the correlation is unknown *a priori* and can be detected only be means of a statistical analysis - seem to be widespread.

### 3.2. Modeling GRN-to-phenotype mapping

As shown above, the GRP map is characterized (except one-to-one mapping) by a significantly larger GRN state space than that of a morphological trait. In terms of Markov model, this means that the construction of a discrete character is a substitution of the original GRN Markov model with a large number of states (Fig. 1a) by a morphological character model with the reduced number of states (Fig. 1b). In this reduced model, at least, some states consist of an aggregated set of the original GRN states. The aggregated states can be unbiasedly modeled if and only if the generating GRN process is lumpable (see Lumpability of Markov models). The conditions for strong lumpability and nearly lumpable chains are likely to be violated in real-life situations due to the rather strict constraints imposed on rate matrix and initial vector – one should not expect that evolution will specifically follow these conditions. The violation can be caused by the complexity of GRN space that will make the aggregation over the GRN states mathematically invalid; so, the aggregated chain will not be lumpable and will preclude the unbiased inference of a morphological character using traditional MM. The bias can be avoided in the HMM context by treating the hidden and observable layers as GRN and trait states respectively (see Handling non-lumpable models). Previous work (Beaulieu et al. 2013) has shown that using MM instead of HMM can bias rate estimates, ancestral state reconstructions and interpretations of rate shifts. Note that hidden states in HMM, besides referring to GRN modules, can be interpreted in other different ways (see Interpreting hidden states). The violation of lumpability and hence biased inference of model parameters is often a result of the character construction procedure. A notable case of this, to which I refer to as the “two-scientist paradox”, is given below.

### 3.3. The two-scientist paradox: hidden Markov models

#### The two-scientist paradox

Suppose two independently evolving genes produce organisms with two traits: (i) red or blue color, and (ii) triangular or round shapes (Fig. 9a-b). Two scientists are willing to reconstruct ancestral states of those traits on a known phylogeny using a model that best fit the data; but the scientists are unfamiliar with the gene-to-trait mapping and each other’s work. The first scientist codes the traits using three different characters: (1) color, (2) shape, and (3) color and shape simultaneously (Fig. 9c). The second scientist scores only one binary character -- presence or absence of being triangular and red simultaneously -- because she or he considers red triangular trait to be a “key innovation” that is worth special attention with respect to all other traits (Fig. 9d). Even though the scientists have different opinions of how to code the traits, they test similar models – traditional MM and HMM – to identify the best one. To demonstrate the consequences of the scientists’ approaches, I generated 100 random trees and characters corresponding to the respective scoring used and ran ancestral character state reconstruction using *corHMM* (Beaulieu et al. 2013); the fit of the models was compared using ΔAIC (Akaike 1974), calculated as AIC_(MM)_-AIC_(HMM)_ (see the scripts and online Appendix S11); the threshold for ΔAIC was interpreted according to Burnham and Anderson (2003).

**Figure 9.**
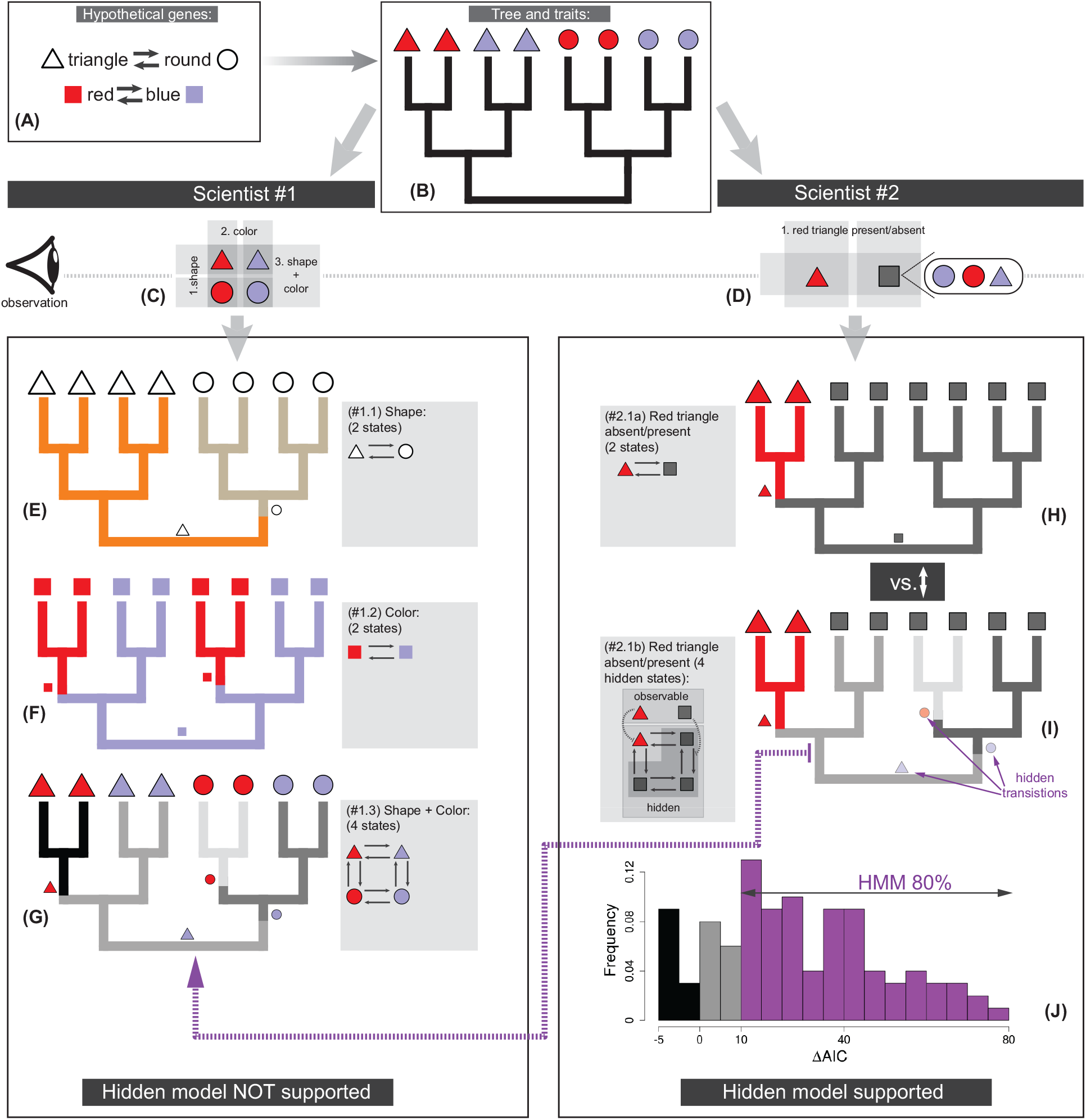
Two-scientist paradox. (A) Two independent genes, that produce color and shape traits, evolve on the phylogeny (B). (C-G) The analyses of the first scientist; neither of the tested models support HMM (only MM models are shown). (D-G) The analyses of the second scientist; HMM used in (I) outperforms MM used in (H). The dashed line from (I) to (G) indicates that HMM in (I) approximates MM of (G). (J) The distribution of ΔAIC values calculated as AIC_(MM, from (H))_-AIC_(HMM, from (I))_; the color on the histogram marks ΔAIC values. The ancestral character state reconstructions in (E-G, H-I) show possible scenario of state evolution that is provided for the explanatory purpose of the two-scientist paradox.

The first scientist finds that traditional MMs have a significantly better fit for all characters, which is expected given the initial conditions for how the traits evolve (Fig. 9e-g). In contrast, the second scientist finds that HMM is favored – it yields a substantially better fit than MM (ΔAIC >10) in 80% of the simulations (Fig. 9h-j). At first glance, this result may seem odd since the initial characters are independent; however the underperformance of MM occurs because the coding scheme of the second scientist corresponds to the partial mapping between the genes and trait meaning that the generating process is incompletely identified with MM; the amalgamated rate matrix that defines simultaneous evolution of the shape and color is not lumpable with respect to that coding scheme, thus the hidden factors have to be included in the model that yields better fit for HMM. The lack of lumpability holds even if the transition rates between and within the genes in the generating process are all equal (online Appendix S11). Note that in the cases where MM performed similar or better (20%), the state aggregation was very close forming a lumpable chain due to the stochastic nature of the simulations, thus, favoring MM.

#### Discussion

Even though, the color and shape traits were simulated with traditional MM, the choice of the coding schemes (as in the case of the second scientist) requires using the different evolutionary model, which, at first glance, is counterintuitive as the initial characters are independently evolving entities. Obviously, different discretization schemes may imply different assumptions of trait evolution; thus, phylogenetic analysis is strongly conditional on the discretization scheme used. Does it mean that the discretization scheme of the first scientist is better than that of the second one? I would argue that it does not, since discretization of traits is always a subjective process. An optimal discretization scheme cannot be selected in advance without prior knowledge of a generating process. In fact, this procedure has to be reversed – an optimal model has to be found that can adequately model the chosen discretization scheme. In this regard, the subjectivity associated with the choice of the discretization becomes minimal. For example, MM of the four-state character of the first scientist, and HMM of the second scientist are different but have similar state spaces, and, in fact, they model similar events – the hidden transitions in HMM tend to approximate the observable transitions of the MM (Fig. 9i,g). Thus, both scientists are doing a good job of modeling their characters since they both perform model comparison and use the best-selected models.

#### Inferring hidden states

The topology of hidden space in HMM (i.e., the number of hidden states and transitions between them) can be anything that does not produce a lumpable model. Since the topology is unknown *a priori* and has to be inferred from observed data (i.e., character states at tips), one needs to test different models by manually varying parameters of the hidden topology and selecting the best models using model selection criteria. This approach can be implemented in *corHMM* or *RevBayes*. The method similar to that of Pagel and Meade (2006) that uses reversible jump Markov Chain Monte Carlo to sample different models from their posterior distribution can be implemented in *RevBayes* but this direction requires further research. When testing different HMMs, one should keep in mind that the data must be identical, which can be achieved as described in the tail color problem. Note that the HMM used in the example above has some similarity to a SMM – its rate matrix is structured to reflect independent evolution between the genes. The techniques reviewed for constructing SMM can be used to build hidden topologies of a HMM.

#### Performance of HMM vs. MM

If lumpability is violated but a traditional MM is used instead of HMM, then it may lead to a significant error in rate estimation and inference of ancestral states. The error magnitude depends on the rate values and rate ratios in the original transition matrix. The substantial bias in rate estimation between MM and variable types of HMM were reported for ancestral character state reconstruction with the empirical and simulated datasets (Beaulieu et al. 2013). If lumpability is violated, the phylogenetic inference may be also biased, which was shown for DNA data when four nucleotides were recoded into fewer groups (Vera-Ruiz et al. 2014). At the same time, if lumpability is valid, state aggregation can be used in algorithms to reduce the size of rate matrix which improves computational performance (Davydov et al. 2017). The simulations for the two-scientist paradox, despite showing a substantially better fit for HMM, detected insignificant differences between HMM and MM rate estimates, which converged on the generating rates (mean squared error was 7×10^-4^ and 9×10^-4^ respectively). This indicates that rate underestimation by MM is context-dependent and if rates are the only focus of inference then, in certain cases, a MM can be used as an appropriate estimator. However, if a study aims at inferring both rates and hidden factors then a HMM should be preferred.

It has to be born in mind that the performance of a HMM is strongly data dependent since, on average, HMM uses more free parameters than MM. The study of Beaulieu et al. (2013) sets the lower limit when HMM can be inferred in ancestral character state reconstruction to 60 - 120 taxa. If fewer taxa are used, then MM can provide a sufficient approximation.

### 3.4. Interpreting hidden states

Initially, HMMs were primarily proposed to model heterogeneity of evolutionary rates across time (Tuffley and Steel 1998; Beaulieu et al. 2013). In this respect hidden states refer to different rate categories or may be also interpreted as hidden extrinsic environmental factors. As was shown above, hidden states can indicate states of GRN modules; also they are conditional on a discretization scheme. Therefore, all four components – time-heterogeneity, GRN transitions, environmental factors, and subjectivity in trait discretization – are confounded in a HMM. In this light, interpreting hidden states must be taken with caution. For example, in the two-scientist paradox, despite the same process that generated the traits, the two scientists may come to drastically different conclusions if the results are interpreted from the perspective of rate shifts. While the first scientist may not report something unexpected, the second scientist may claim that the “key innovation” of being triangular and red was subjected to complex shifts during evolution. Such conclusion, depending on the scientist’s imagination, can be connected with some other biological observations to result in a “spectacular” finding, which, in fact, is misleading. In general, I suggest using a moderate interpretation of hidden states – as a simultaneous realization of all confounding processes mentioned above.

## Discussion

The present paper lays out a framework for modeling hierarchical and hidden processes whose realization produce discrete phenotypic traits. While both processes are at first glance dissimilar, they are in fact interacting -- modeling a hierarchical process often requires equipping a SMM with a HMM; at the same time modeling hidden processes requires structuring HMM using SMM techniques. Besides shared math, this occurs because the hierarchical process is partially driven by the hidden process - formation of anatomical dependencies is the result of sequential realizations of GRN modules during embryo development.

The main suggestion of this paper is to use a joint structured Markov model with hidden states (SMM+HMM) that, as was shown, can accommodate multitude processes driving trait evolution. Usually, several alternative models can be proposed to code the same trait; they can be tested by keeping the data (number of hidden states per each observable characters) the same and using model selection procedure. The included simulation studies demonstrate how the proposed framework can be implemented in practice. Additionally, this paper summarizes all main techniques which can be used to amalgamate, decompose and structure morphological characters in a mathematically consistent way, which gives rigorous grounds for coding traits into character. The proposed approaches can be used for developing new methods aiming at reconstructing correlated trait evolution and ancestral ontologies.

Structured Markov models equipped with hidden sates provide a natural way to model anatomical dependencies between traits for phylogenetic analyses. These models give unambiguous solutions to the tail color problem and can be flexibly extended to model complex hierarchies as was shown using the tail armor case. The provided solutions via SMM-ind and SMM-sw can be further generalized using SMM-gen to account for various types of relationships which may exist between traits. The efficient use of SMM requires an identification of anatomical dependencies and their incorporation through the appropriate structuring of rate matrix. This means that trait coding and model selection are the parts of the same analytical procedure as their choice is affected by organismal anatomy. I suggest that hierarchically dependent traits have to be coded as one character despite that SMM-ind can be represented using two characters. First of all, one-character representation will help to avoid data recoding if both models - SMM-ind and SMM-sw - are tested for the same dataset. Second, if SMM-ind is extended to have more free parameters as, for example, SMM-gen or a model similar to F81 (Felsenstein 1981), then its two-character representation will no longer be valid, and this extension will require only one multistate character. For phylogenetic inference, all characters of a dataset that encode a set of interrelated dependencies can be placed in a single data partition subset to which a selected model is assigned. This strategy will allow testing different candidate models using the same data.

The artificial recoding of characters (e.g., when a multistate character is recoded to a set of binary characters) should not be used - it introduces spurious model assumptions that are inconsistent with the properties of character amalgamation and state aggregation. At the same time, the invariance of character and character state can be a helpful tool as it, to a certain extent, eliminates the ambiguity of discretizing traits into character and character states. Trait scoring and model choice must be done thoughtfully and must be consistent with the underlying mathematics, organismal anatomies, and biological knowledge.

Hidden Markov models provide an insight into the underlying hidden process that cannot be achieved through MM. Trait discretization is a strongly subjective procedure but application of an appropriate HMM minimizes this subjectivity. Interpreting state shifts in the phylogenetic context must be taken with caution. Hidden Markov models will constantly overperform MM if the generating process is not lumpable and the amount of data is sufficient. The use of HMM is especially essential for analyzing large datasets which naturally contain heterogeneity. The model selection between MM and HMM must be an obligatory component of, at least, ancestral character state reconstruction since even a complex MM (because of lumpability violations) might be inappropriate for inference. The type of HMM discussed in the present study can be also used in tree inference with morphological data, however, HMM application will be limited due to stricter model parameterization. Such applications will likely require the assignment of the same HMM with *a priori* specified hidden states to all characters of a partition subset.

In phylogenetics, every character statement is a hypothesis of primary homology (Hawkins et al. 1997). The “real” assessment of whether an observation in one species is homologous or homoplasious with respect to that in the other can be done only through a phylogenetic inference or reconstruction of character evolution (secondary homology). It has been considered that a thoughtful identification of primary homology is a prerequisite for a successful phylogenetic analysis. Obviously, in traditional MM and parsimony, the secondary homology is conditional on primary homology (Agnarsson and Coddington 2008; Brazeau 2011). The discordance between primary and secondary homology, to a large extent, occurs due to the discordance between the morphological traits and underlying GRN states. Hidden Markov models are capable of directly accounting for this discordance. Theoretically, this means that the quality of primary homology statement should not matter if HMM is used since it can automatically adjust the underlying hidden space. The practical aspect of this consideration requires further research though.

## Supporting information

## Supplementary Materials

Online Appendix and Supplementary materials are available at bioRxiv.

## Acknowledgements

Thanks are due to Josef Uyeda (Virginia Tech), Sergey Gavrilets (University of Tennessee), Sarah Flanagan (National Institute for Mathematical and Biological Synthesis, University of Tennessee), Brian O’Meara (University of Tennessee), Dimitar Dimitrov (University of Bergen), Martín Ramírez (Museo Argentino de Ciencias Naturales), Emma Goldberg (University of Minnesota) and six anonymous reviewers for useful suggestions on the text of the manuscript.

