## Supplementary material for "Integration of anatomy ontologies and evo-devo using structured Markov models suggests a new framework for modeling discrete phenotypic traits"

SERGEI TARASOV

#### S1. Two characters: independent evolution

Suppose there are two initial two-state characters: ***X*** {$x_1$, $x_2$} and ***Y*** {$y_1$, $y_2$} which we wish to amalgamate into one single character ***Z***. The initial characters ***X*** and ***Y*** are defined as:

$$\boldsymbol{X} = \begin{matrix} & \boldsymbol{x_1} & \boldsymbol{x_2} \\ \boldsymbol{x_1} & -\alpha & \alpha \\ \boldsymbol{x_2} & \beta & -\beta \end{matrix};\ \boldsymbol{Y} = \begin{matrix} & \boldsymbol{y_1} & \boldsymbol{y_2} \\ \boldsymbol{y_1} & -\gamma & \gamma \\ \boldsymbol{y_2} & \theta & -\theta \end{matrix}.$$

The matrices of transition probabilities (i.e., $P(t)$) for ***X*** and ***Y*** after some time *t* elapsed are:

$$\boldsymbol{P_X}(t) = e^{\boldsymbol{X}t} \text{ and } \boldsymbol{P_Y}(t) = e^{\boldsymbol{Y}t}.$$

The brute way to get the state probabilities of ***Z*** (i.e., $\boldsymbol{P_z}(t)$) is to manually multiply every single element of matrix $\boldsymbol{P_X}(\boldsymbol{t})$ by every single element of matrix $\boldsymbol{P_Y}(\boldsymbol{t})$. This will yield a four-by-four transition matrix. The four states of $\boldsymbol{P_z}(t)$ correspond to all possible combinations of states between ***X*** and ***Y***, which are {$x_1y_1$, $x_2y_1$, $x_1y_2$, $x_2y_2$}. The same operation can be done analytically using the Kronecker product:

$$\boldsymbol{P_z}(t) = \boldsymbol{P_X}(t) \otimes \boldsymbol{P_Y}(t).$$

There is a connection between matrix exponentiation and the Kronecker product [*Theorem 1*, in Steeba and Wilhelm (1981)] that allows expressing the transition probability matrix of the combined chain using the initial rate matrices:

$$\boldsymbol{P_z}(t) = \boldsymbol{P_X}(t) \otimes \boldsymbol{P_Y}(t) = e^{(\boldsymbol{X} \otimes \boldsymbol{I_Y} + \boldsymbol{I_X} \otimes \boldsymbol{Y})t}, \text{ (S1.1)}$$

where $\boldsymbol{I_Y}$ and $\boldsymbol{I_X}$ are identity matrices of the same dimension as matrix ***Y*** and ***X*** respectively (i.e., dimension=2). The right hand side of the equation (S1.1) suggests that the rate matrix ***Z*** is:

$$\boldsymbol{Z} = \boldsymbol{X} \otimes \boldsymbol{I_Y} + \boldsymbol{I_X} \otimes \boldsymbol{Y}. \text{ (S1.2)}$$

Given the equation (S1.2), the matrix ***Z*** is:

$$\boldsymbol{Z} = \begin{matrix} \\ \boldsymbol{x_1y_1} \\ \boldsymbol{x_1y_2} \\ \boldsymbol{x_2y_1} \\ \boldsymbol{x_2y_2} \end{matrix} \begin{matrix} \boldsymbol{x_1y_1} & \boldsymbol{x_1y_2} & \boldsymbol{x_2y_1} & \boldsymbol{x_2y_2} \\ \end{matrix} \begin{pmatrix} -\alpha-\gamma & \gamma & \alpha & 0 \\ \theta & -\alpha-\theta & 0 & \alpha \\ \beta & 0 & -\beta-\gamma & \gamma \\ 0 & \beta & \theta & -\beta-\theta \end{pmatrix}. \text{ (S1.3)}$$

#### S2. *Character and character state invariance.*

The tree inference was run in *RevBayes* (Höhna et al. 2016) for the two datasets: (i) three two-state characters (Fig. 2a), and (ii) one four-state character (Fig. 2b). The datasets were constructed by replicating character (i) 30 times and character (ii) 10 times (the amalgamation of three characters gives one character). The replication was done to make data sufficiently informative for the inference. The dataset of character (i) was analyzed using two-state *Mk* model (S2.1), while the dataset from the character (ii) using *Mk* model with four states (S2.2), and structured *MK* model (*Mk-SMM-ind*). The amalgamation of three two-state characters (i) results in eight-state *Mk-SMM-ind* (S2.3); four of these states *{001, 010, 011, 101}* are not observed in the provided set of species (Fig. 2) and they were removed that resulted in the final four-state rate matrix (S2.4) used in inference. This was done as, at the moment, *RevBayes* cannot handle states given in rate matrix but not scored for the observed species. If the functionality of using unobserved states in the rate matrix is implemented, the use of matrix (S2.3) is possible. The analyses were run for $10^6$ generations, using the exponential prior for branch lengths ~ *Exp(0.2)*.

$$Mk = \begin{pmatrix} \mathbf{0} & \mathbf{1} \\ - & 1 \\ 1 & - \end{pmatrix}, \text{ (S2.1)}$$

$$Mk = \begin{pmatrix} \mathbf{0} & \mathbf{1} & \mathbf{2} & \mathbf{3} \\ - & 1 & 1 & 1 \\ 1 & - & 1 & 1 \\ 1 & 1 & - & 1 \\ 1 & 1 & 1 & - \end{pmatrix}, \text{ (S2.2)}$$

$$Mk-SMM-ind = \begin{pmatrix} \mathbf{000} & \mathbf{001} & \mathbf{010} & \mathbf{011} & \mathbf{100} & \mathbf{101} & \mathbf{110} & \mathbf{111} \\ - & 1 & 1 & 0 & 1 & 0 & 0 & 0 \\ 1 & - & 0 & 1 & 0 & 1 & 0 & 0 \\ 1 & 0 & - & 1 & 0 & 0 & 1 & 0 \\ 0 & 1 & 1 & - & 0 & 0 & 0 & 1 \\ 1 & 0 & 0 & 0 & - & 1 & 1 & 0 \\ 0 & 1 & 0 & 0 & 1 & - & 0 & 1 \\ 0 & 0 & 1 & 0 & 1 & 0 & - & 1 \\ 0 & 0 & 0 & 1 & 0 & 1 & 1 & - \end{pmatrix}, \text{ (S2.3)}$$

$$Mk-SMM-ind = \begin{pmatrix} \mathbf{0} & \mathbf{1} & \mathbf{2} & \mathbf{3} \\ - & 1 & 0 & 0 \\ 1 & - & 1 & 0 \\ 0 & 1 & - & 1 \\ 0 & 0 & 1 & - \end{pmatrix}. \text{ (S2.4)}$$

#### S3a. Character state aggregation

The character states of $m \times m$ rate matrix $\boldsymbol{M}$ can be aggregated (if the matrix is lumpable) given the partitioning scheme $\mathcal{B}$ to result in $n \times n$ matrix $\boldsymbol{M'}$:

$$\boldsymbol{M'} = A_{ggr.}(\boldsymbol{M}|\mathcal{B}) = \mathbf{N}\mathbf{P}\boldsymbol{M}\mathbf{P}^{\mathrm{T}}$$

where **N** is a $n \times n$ diagonal matrix with $1/s$ elements ($s$ is the number of the initial states in the aggregated state), **P** is a $n \times m$ matrix with 1's indicating assignment of the initial state to a specified aggregated state, and superscript **T** indicates matrix transpose.

The amalgamated tail-color character $A_{mal.(ind)}(\boldsymbol{T},\boldsymbol{C})$ from the equation (2) can be split under the two partitioning schemes $\mathcal{B}_1 = \{\{ar, ab\}, \{pr, pb\}\}$ and $\mathcal{B}_2 = \{\{ar, pr\}, \{ab, pb\}\}$:

$$A_{mal.(ind)}(\boldsymbol{T},\boldsymbol{C}) = \boldsymbol{M} = \begin{pmatrix} \mathbf{00} & \mathbf{01} & \mathbf{10} & \mathbf{11} \\ -\alpha-\gamma & \gamma & \alpha & 0 \\ \theta & -\alpha-\theta & 0 & \alpha \\ \beta & 0 & -\beta-\gamma & \gamma \\ 0 & \beta & \theta & -\beta-\theta \end{pmatrix},$$

$$A_{ggr.}(\boldsymbol{M}|\mathcal{B}_1) = \begin{pmatrix} \frac{1}{2} & 0 \\ 0 & \frac{1}{2} \end{pmatrix} \begin{pmatrix} 1 & 1 & 0 & 0 \\ 0 & 0 & 1 & 1 \end{pmatrix} \boldsymbol{M} \begin{pmatrix} 1 & 1 & 0 & 0 \\ 0 & 0 & 1 & 1 \end{pmatrix}^{\mathrm{T}} = \begin{pmatrix} \mathbf{0} & \mathbf{1} \\ -\alpha & \alpha \\ \beta & -\beta \end{pmatrix},$$

$$A_{ggr.}(\boldsymbol{M}|\mathcal{B}_2) = \begin{pmatrix} \frac{1}{2} & 0 \\ 0 & \frac{1}{2} \end{pmatrix} \begin{pmatrix} 1 & 0 & 1 & 0 \\ 0 & 1 & 0 & 1 \end{pmatrix} \boldsymbol{M} \begin{pmatrix} 1 & 0 & 1 & 0 \\ 0 & 1 & 0 & 1 \end{pmatrix}^{\mathrm{T}} = \begin{pmatrix} \mathbf{0} & \mathbf{1} \\ -\gamma & \gamma \\ \theta & -\theta \end{pmatrix}.$$

#### S3b. Two characters: general case of correlation

As in the section (S1), suppose there are two initial characters $\boldsymbol{X}$ and $\boldsymbol{Y}$ which we wish to amalgamate into a single character $\boldsymbol{Z}$ under the condition that the characters $\boldsymbol{X}, \boldsymbol{Y}$ are fully correlated. Unlike the matrix of independent evolution (S1.3), that is characterized by the four rate parameters, the general case of correlation requires eight rate parameters and thereby cannot be derived from the two initial matrices. So, the combined rate matrix **Z** has to be characterized by four initial matrices $\boldsymbol{X_1}$, $\boldsymbol{X_2}$ and $\boldsymbol{Y_1}$, $\boldsymbol{Y_2}$ corresponding to the characters $\boldsymbol{X}$ and $\boldsymbol{Y}$ respectively:

$$\boldsymbol{X_1} = \begin{pmatrix} -\alpha & \alpha \\ \beta & -\beta \end{pmatrix}, \boldsymbol{Y_1} = \begin{pmatrix} -\gamma & \gamma \\ \theta & -\theta \end{pmatrix}, \boldsymbol{X_2} = \begin{pmatrix} -\alpha_1 & \alpha_1 \\ \beta_1 & -\beta_1 \end{pmatrix}, \boldsymbol{Y_2} = \begin{pmatrix} -\gamma_1 & \gamma_1 \\ \theta_1 & -\theta_1 \end{pmatrix}.$$

The interdependencies between the rates in these matrices can be interpreted as follows:

$\boldsymbol{X_1}$ depends on $\boldsymbol{Y_1}$ state *1*; and $\boldsymbol{Y_1}$ depends on $\boldsymbol{X_1}$ state *2*;

$\boldsymbol{X_2}$ depends on $\boldsymbol{Y_2}$ state *2*; and $\boldsymbol{Y_2}$ depends on $\boldsymbol{X_2}$ state *1*.

The two-step procedure is needed to get $\boldsymbol{Z}$: (1) obtaining correlated matrix for each pair $\boldsymbol{X_1}$, $\boldsymbol{Y_1}$ and $\boldsymbol{X_2}$, $\boldsymbol{Y_2}$ using a method similar to the previous example, and then (2) adding up these two correlated matrices into the final one:

$$\boldsymbol{Z} = [\boldsymbol{X_1} \otimes \begin{pmatrix} 1 & 0 \\ 0 & 0 \end{pmatrix} + \begin{pmatrix} 0 & 0 \\ 0 & 1 \end{pmatrix} \otimes \boldsymbol{Y_1}] + [\boldsymbol{X_2} \otimes \begin{pmatrix} 0 & 0 \\ 0 & 1 \end{pmatrix} + \begin{pmatrix} 1 & 0 \\ 0 & 0 \end{pmatrix} \otimes \boldsymbol{Y_2}].$$

This equation gives:

$$\boldsymbol{Z} = \begin{pmatrix} -\alpha - \gamma_1 & \gamma_1 & \alpha & 0 \\ \theta_1 & -\alpha_1 - \theta_1 & 0 & \alpha_1 \\ \beta & 0 & -\beta - \gamma & \gamma \\ 0 & \beta_1 & \theta & -\theta - \beta_1 \end{pmatrix}.$$

#### S3c. Two characters: "switch-on" case of correlation

Suppose that during the course of evolution both states of character *Y* can appear only if the character *X* is in the state $\boldsymbol{x_2}$; if X is in the state $\boldsymbol{x_1}$ then the character *Y* is "switched-off". The modified version of the equation (S1.2) has to be used to construct the amalgamated matrix **Z**. This modification assigns zero value in the identity matrix $\boldsymbol{I_X}$ to the state which does not imply dependency (i.e. $\boldsymbol{x_1}$) and leave value one for the state which implies it (i.e. $\boldsymbol{x_2}$). The modified $\boldsymbol{I_X}$ becomes the diagonal matrix $\boldsymbol{D_X}$. In matrix notation this leads to the following:

$$\boldsymbol{I_X} = \begin{pmatrix} 1 & 0 \\ 0 & 1 \end{pmatrix} \rightarrow \boldsymbol{D_X} = \begin{pmatrix} 0 & 0 \\ 0 & 1 \end{pmatrix}.$$

Next, the matrix $\boldsymbol{D_X}$ can be plugged in the equation (S1.2) instead of $\boldsymbol{I_X}$, to get the rate matrix of the correlated character, which yields:

$$\boldsymbol{Z} = \boldsymbol{X} \otimes \boldsymbol{I_Y} + \boldsymbol{D_X} \otimes \boldsymbol{Y} = \begin{pmatrix} -\alpha & 0 & \alpha & 0 \\ 0 & -\alpha & 0 & \alpha \\ \beta & 0 & -\beta - \gamma & \gamma \\ 0 & \beta & \theta & -\beta - \theta \end{pmatrix}.$$

#### Table S1. The coding schemes for the tail color problem.

| Scheme | Species | Tail presence | Tail color | | Amalgamated state space |
|---|---|---|---|---|---|
| **#1** | **No tail** | 0 | ? *(means 0 or 1)* | | {00, 01, 10, 11} |
| | **Red** | 1 | 0 | | |
| | **Blue** | 1 | 1 | | |
| **#2** | | **Tail presence** | **Tail color** | | {00, 01, 02, 10, 11, 12} |
| | **No tail** | 0 | 0 | | |
| | **Red** | 1 | 1 | | |
| | **Blue** | 1 | 2 | | |
| **#3** | | **Tail presence** | **Red tail presence** | **Blue tail presence** | {000, 001, 010, 011, 100, 101, 110, 111} |
| | **No tail** | 0 | 0 | 0 | |
| | **Red** | 1 | 1 | 0 | |
| | **Blue** | 1 | 0 | 1 | |
| **#4** | | **Tail** | | | N/A, {0, 1, 2} |
| | **No tail** | 0 | | | |
| | **Red** | 1 | | | |
| | **Blue** | 2 | | | |
| **SMM** | | **Tail** | | | N/A, {0, 1, 2, 3} |
| | **No tail** | 0&1 *(means 0 or 1)* | | | |
| | **Red** | 2 | | | |
| | **Blue** | 3 | | | |

#### S4. Weak lumpability and equilibrium distribution

Interesting case of weak lumpability arises when the initial vector of Markov chain coincides with the equilibrium distribution of that chain. Such chain is weakly lumpable in respect to any possible partitioning scheme. Consider four-state $\{s_1, s_2, s_3, s_4\}$ rate matrix $Q$. Suppose its initial vector $\pi_{st}$ has equilibrium probability that implies:

$$\pi_{st}Q = 0 \quad \text{(S4.1)}$$

The probability of states in this chain after some time is time-invariant, i.e., $P(t)=\pi_{st}e^{Qt}=\pi_{st}$. If we aggregate this chain under the partitioning $\mathcal{B}=\{\{s_1, s_2\}, \{s_3, s_4\}\}$, then the aggregated initial vector becomes $A_{ggr.}(\pi_{st})=\{\{\pi_{st1}+\pi_{st2},\}, \{\pi_{st3}+\pi_{st3}\}\}$; there exists a two-state stationary Markov chain defined by the rate matrix $A_{ggr.}(Q)$ whose initial vector with equilibrium probabilities is $A_{ggr.}(\pi_{st})$, so the original chain is weakly lumpable, which can be expressed as:

$$A_{ggr.}(\pi_{st}Q) = A_{ggr.}(\pi_{st})A_{ggr.}(Q) = 0 \quad \text{(S4.2)}$$

In fact, there is a whole family of such two-state chains that can be defined by different rate matrices but have the same equilibrium probabilities. Moreover, such weak lumpability would hold for any possible partitioning scheme of $Q$. Generally, weak lumpability requires some notable symmetries in the initial vector, as for example equilibrium distribution or equal probabilities (Rubino and Sericola 1993, 2014); thus, weak lumpability holds only under the strict constraints of the initial vector.

Weak lumpability is an attractive property but it does not seem to work in phylogenetic analysis due to the way likelihood is calculated on trees; this calculation does not preserve the lumpability of the model. I will demonstrate it by showing a simple example where the model is weakly lumpable but not strongly lumpable. To derive this, without loss of generality, I use a simple discrete-time Markov process that is easier to work with than the continuous-time process. The equality given by the equation S4.2 is the same for the discrete-time process except that it does not equal to 0.

Consider a phylogenetic tree that consists of two tips $t_1$ and $t_2$ descending from the same node $n_1$. Each tip is a column vector summarizing observations (i.e. character); the elements of the vectors $t_1$ and $t_2$ are set to 1 if the character state is observed and zero otherwise. Suppose that we are dealing with a three-state character and both $t_1$ and $t_2$ are the same $t_1$= $t_2$= {0,0,1}. Define $L(n_1)$ to be the conditional probability of data descending from $n_1$; suppose that we calculate $L(n_1)$ given that all branch lengths are equal to 1, and the discrete-time transition matrix $Q$ is:

$$Q=\begin{pmatrix} 1-2a & a & a \\ a & 1-a-b & b \\ a & b & 1-a-b \end{pmatrix}. \text{ (S4.3)}$$

The initial probability vector has equilibrium distribution that is $\pi_{st} = \{\frac{1}{3}, \frac{1}{3}, \frac{1}{3}\}$. Suppose, we lump this matrix given the partitioning scheme $\mathcal{B}$ ={{1, 2}, {3}}, so the process is weakly lumpable but not strongly lumpable under this partitioning scheme. The aggregated initial vector becomes $A_{ggr.}(\pi_{st}) = \{\frac{2}{3}, \frac{1}{3}\}$. Let's derive the transition matrix $A_{ggr.}(Q)$ for this lumped process. Aggregating the transitions *3->1* and *3->2* is straightforward since the lumped transition is the sum of the initial ones: *a+b*; the aggregation of the transitions *1->3* and *2->3* can be derived by considering $A_{ggr.}(\pi_{st})$. The only value of *{1,2}->3* that preserves equality $A_{ggr.}(\pi_{st})A_{ggr.}(\boldsymbol{Q}) = A_{ggr.}(\pi_{st})$ is $\frac{a+b}{2}$. So the lumped matrix is:

$$A_{ggr.}(\boldsymbol{Q}) = \begin{pmatrix} 1-\frac{a+b}{2} & \frac{a+b}{2} \\ a+b & 1-a-b \end{pmatrix}. \text{ (S4.4)}$$

The conditional probability for node $n_1$ using the initial process is:

$$L(n_1) = \pi_{st}(Qt_1)(Qt_2). \text{ (S4.5)}$$

The same using the lumped process:

$$L_{aggr.}(n_1) = A_{ggr.}(\pi_{st})(A_{ggr.}(Q)t_1)(A_{ggr.}(Q)t_2). \text{ (S4.6)}$$

If lumpability property is preserved during likelihood calculation than the following equality must hold:

$$A_{ggr.}(L(n_1)) = L_{aggr.}(n_1). \text{ (S4.6)}$$

Given that $t_1$ and $t_2$ are the same, the conditional probabilities are:

$$L(n_1) = (\frac{1}{3}, \frac{1}{3}, \frac{1}{3}) \begin{pmatrix} a \\ b \\ -a-b+1 \end{pmatrix}^2 \rightarrow A_{ggr.}(L(n_1)) = \begin{pmatrix} \frac{1}{3}(a^2+b^2) \\ \frac{1}{3}(-a-b+1)^2 \end{pmatrix}$$

$$L_{aggr.}(n_1) = (\frac{2}{3}, \frac{1}{3}) \begin{pmatrix} \frac{a+b}{2} \\ -a-b+1 \end{pmatrix}^2 = \begin{pmatrix} \frac{1}{6}(a+b)^2 \\ \frac{1}{3}(-a-b+1)^2 \end{pmatrix}$$

Apparently $A_{ggr.}(L(n_1)) \neq L_{aggr.}(n_1)$ because their first elements are different; the summing over the elements of $A_{ggr.}(L(n_1))$ and $L_{aggr.}(n_1)$ will, likewise, result in different likelihood values.

Despite that the model under the consideration is weakly lumpable, this lumpability does not hold for the likelihood calculation due to the way the multiplications are carried out in the pruning algorithm. Note that if *a=b* in the initial matrix *Q* (so the model is strongly lumpable), then the equality S8.6 holds.

#### S5. Nearly lumpable chains

In this paper, the term nearly lumpable Markov model refers to such Markov model that violates the conditions of strong lumpability but if aggregated can be used to approximate the behavior of the original model with insignificant error. I demonstrate the property of nearly lumpable model by aggregating states in the original model $X^*$ and thereby constructing the aggregated model $Y^*$.

*The Markov chain $X^*$ is a continuous-time Markov chain, defining correlated evolution of a sufficiently large number of initial Markov chains; it is nearly lumpable if the following conditions hold: (1) values of its rate matrix are i.i.d. random variables; (2) values of the initial vector $\pi$ are i.i.d. random variables; (3) each state of the aggregated chain $Y^*$ is composed of a sufficiently large number of states of $X^*$.*

In derivation below, I substitute, without loss of generality, the continuous-time Markov chain $X^*$ and $Y^*$ by a discrete-time Markov chains *X* and *Y*. This transformation is done by the uniformization method (Stewart 2009) that is usually used for analyzing continuous-time Markov chains. The obtained discrete-time Markov chain preserves the matrix symmetries and *i.i.d.* property of the matrix entities of $X^*$. The matrix entities of the discrete-time chain *X* are no longer rates but probabilities.

Suppose there is an original discrete-time Markov chain *X* with state space $S = \{1, 2 \ldots n_s\}$ describing fully correlated evolution of a large number of elementary characters. The chain *X* is defined by the initial probability vector $\pi$ and transition matrix **Q** whose elements are probabilities $q_{ij}$ specifying transitions from state $S_i$ to state $S_j$ in *X*. The probabilities in **Q** are assumed to be identically and independently distributed (*i.i.d*) random variables having the same probability distribution $f_q$. The elements $\pi_i$ of the initial vector $\pi$ are also *i.i.d.* random variables having the same probability distribution $f_\pi$.

The chain $Y$=*agg*($\pi$, **Q**, B, $X$) is a function of $X$; it is constructed by aggregating the state space of $X$ into $M$ proper subsets given the partitioning scheme $\mathcal{B}$ ={$B$(1) … $B$(M)}; the state space of $Y$ is denoted by $F$ = {1 … M}. Each element of $F$, denoted as $F_k$ is composed of some states $S$ of $X$; the number of states of $S$ that are contained in $F_k$ is denoted as $N(F_k)$. The chain $Y$ is defined by $\widehat{\mathbf{Q}}$ and $\hat{\pi}$ referring to the transition matrix, and initial probability vector respectively. Matrix $\widehat{\mathbf{Q}}$ consists of elements $\hat{q}_{k,l}$ which are the probabilities of transition from some state $F_k$ to $F_l$. In matrix notations, the aggregation of states is a grouping of cells $q_{i,j}$ of **Q** given $\mathcal{B}$ into the matrix blocks of dimension $N(F_k) \times N(F_l)$ corresponding to the transition rates between state $F_k$ and $F_l$ in $Y$; these matrix blocks are denoted as $\tilde{F}_{k,l}$.

Let us consider any $\tilde{F}_{k,l}$ and let us denote by $\varphi_i$ the sum of $q_{i,j}$ elements in row $i$ of $\tilde{F}_{k,l}$:

$$\varphi_i = \sum_{q_{i,j} \epsilon \tilde{F}_{k,l}} q_{i,j}. \text{ (S5.1)}$$

The strong lumpability specifies that the chain $X$ is lumpable given the partitioning scheme $\mathcal{B}$, if all $\varphi_i$ of $\tilde{F}_{k,l}$ are equal and this holds for all $\tilde{F}_{k,l}$. In this case the probability $\hat{q}_{k,l}$ of the aggregated chain is any $\varphi_i$ of $\tilde{F}_{k,l}$. The aggregated initial vector $\hat{\pi}_k$ corresponding to $\tilde{F}_{k,l}$ is constructed by summing over all $\pi_i$ of $\tilde{F}_{k,l}$:

$$\hat{\pi}_k = \sum_{\pi_i \epsilon \tilde{F}_{k,l}} \pi_i,$$

The strong lumpability implies the two following statements which guaranty the lumpability:

$$\hat{q}_{k,l} = \frac{\sum_{\varphi_i \epsilon \tilde{F}_{k,l}} \varphi_i}{N(F_k)}, \text{ (S5.2)}$$

$$\sum_{\varphi_i \epsilon \tilde{F}_{k,l}} \pi_i \varphi_i = \pi_k \hat{q}_{k,l}. \text{ (S5.3)}$$

Let us express RHS of the equation (S5.3) using the equation (S5.2):

$$\sum_{\varphi_i \epsilon \tilde{F}_{k,l}} \pi_i \varphi_i = \pi_k \frac{\sum_{\varphi_i \epsilon \tilde{F}_{k,l}} \varphi_i}{N(F_k)}. \text{ (S5.4)}$$

Below, I demonstrate that LHS and RHS of the equation (S5.4) converge when the chain $X$ is aggregated given the condition specified above. This convergence maintains the lumpability of $X$. In deriving this convergence, I suggest that $\varphi_i$ of $\tilde{F}_{k,l}$ is *i.i.d.* random variable coming from some probability distribution. This suggestion is based on two facts: (i) $q_{i,j}$ is *i.i.d.* by the initial conditions, (ii) rows of $\tilde{F}_{k,l}$ are randomly visited by Markov chain given the random nature of the initial vector.

The LHS of the equation (S5.4) can be re-written in the following way:

$$\sum_{\varphi_i \epsilon \tilde{F}_{k,l}} \pi_i \varphi_i = \frac{\sum_{\varphi_i \epsilon \tilde{F}_{k,l}} \pi_i \varphi_i}{N(F_k)} N(F_k), \text{ (S5.5)}$$

where $\frac{\sum_{\varphi_i \epsilon \tilde{F}_{k,l}} \pi_i \varphi_i}{N(F_k)}$ is a mean value of the product of two random variables $\pi_i$ and $\varphi_i$. Let's denote it by $\mu(\pi_i \varphi_i)$. As $N(F_k)$ increases, $\mu(\pi_i \varphi_i)$ converges to its expectation $E(\pi_i \varphi_i)$; the expectation of a product of random variables is the product of their expectations, which if written using their means gives:

$$E(\pi_i \varphi_i) = \mu(\varphi_i)\mu(\pi_i).\ \text{(S5.6)}$$

Now, the RHS of the equation (S5.5) can be expressed as:

$$\frac{\sum_{\varphi_i \epsilon \tilde{F}_{k,l}} \pi_i \varphi_i}{N(F_k)} N(F_k) = E(\pi_i \varphi_i) N(F_k) = \mu(\varphi_i)\mu(\pi_i)N(F_k),\ \text{(S5.7)}$$

where, $\mu(\varphi_i) = \frac{\sum_{\varphi_i \epsilon \tilde{F}_{k,l}} \varphi_i}{N(F_k)}$ and $\mu(\pi_i) = \frac{\pi_k}{N(F_k)}$. So, the RHS of the equation (S5.7) gives:

$$\mu(\varphi_i)\mu(\pi_i)N(F_k) = \frac{\sum_{\varphi_i \epsilon \tilde{F}_{k,l}} \varphi_i}{N(F_k)} \frac{\pi_k}{N(F_k)} N(F_k) = \pi_k \frac{\sum_{\varphi_i \epsilon \tilde{F}_{k,l}} \varphi_i}{N(F_k)}.\ \text{(S5.8)}$$

The equation (S5.8) is derived from the LHS of the equation (S5.4) suggesting that the LHS and RHS of the equation (S5.4) converge. The initial conditions allow extrapolating this convergence to all $\tilde{F}_{k,l}$ induced by the partitioning scheme. So, the original chain *X* is nearly lumpable.

#### S6. Lumping SMM-ind and SMM-sw

The SMM-ind and SMM-sw matrices for the scheme #1 cannot be amalgamated into a three state matrix with states {*a,r,b*}. This can be verified by checking that the following equality necessary for lumpability does not hold under the partitioning {{*ar, ab*}, {*pr*}, {*pb*}}:

$$\mathbf{P}^{\mathbf{T}}\mathbf{NP}\boldsymbol{M}\mathbf{P}^{\mathbf{T}} = \boldsymbol{M}\mathbf{P}^{\mathbf{T}},\ \text{(S6.1)}$$

where ***M*** is either SMM-ind or SMM-sw matrix respectively. This happens because in both SMM-ind and SMM-sw the following rates are different: *ab*→*pb* ≠ *ar*→*pb*, and *ab*→*pr* ≠ *ar*→*pr*.

The SMM-sw for the scheme #1 cannot be lumped to yield an independent tail color character that implies lumping the SMM-sw under the partitioning scheme $B_2 = \{\{ar, pr\}, \{ab, pb\}\}$. This can be verified using the equation (S6.1).

#### S7. Mk-like models for tail color problem

Below, I provide *Mk*-like parametrization for SMM-ind and SMM-sw. Noteworthy, that these matrices are not scaled to express branch lengths in the expected number of changes per site. To scale them in RevBayes, use the function *fnFreeK(rescaled=true, …).*

$$\textit{Mk-SMM-ind} = \begin{pmatrix} \mathbf{00} & \mathbf{01} & \mathbf{10} & \mathbf{11} \\ - & 1 & 1 & 0 \\ 1 & - & 0 & 1 \\ 1 & 0 & - & 1 \\ 0 & 1 & 1 & - \end{pmatrix}$$

$$\textit{Mk-SMM-sw} = \begin{pmatrix} \mathbf{00} & \mathbf{01} & \mathbf{10} & \mathbf{11} \\ - & 0 & 1 & 0 \\ 0 & - & 0 & 1 \\ 1 & 0 & - & 1 \\ 0 & 1 & 1 & - \end{pmatrix}$$

#### S8. Mk-like models for tail armor case

Below, I provide *Mk*-like parametrization for SMM-ind and SMM-sw. Noteworthy, that these matrices are not scaled to express branch lengths in the expected number of changes per site. To scale them in RevBayes, use the function *fnFreeK(rescaled=true, …).*

$$Mk-SMM-ind = \begin{pmatrix} \mathbf{000} & \mathbf{001} & \mathbf{010} & \mathbf{011} & \mathbf{100} & \mathbf{101} & \mathbf{110} & \mathbf{111} \\ - & 1 & 1 & 0 & 1 & 0 & 0 & 0 \\ 1 & - & 0 & 1 & 0 & 1 & 0 & 0 \\ 1 & 0 & - & 1 & 0 & 0 & 1 & 0 \\ 0 & 1 & 1 & - & 0 & 0 & 0 & 1 \\ 1 & 0 & 0 & 0 & - & 1 & 1 & 0 \\ 0 & 1 & 0 & 0 & 1 & - & 0 & 1 \\ 0 & 0 & 1 & 0 & 1 & 0 & - & 1 \\ 0 & 0 & 0 & 1 & 0 & 1 & 1 & - \end{pmatrix}$$

$$Mk\text{-}SMM\text{-}sw = \begin{pmatrix} \mathbf{000} & \mathbf{001} & \mathbf{010} & \mathbf{011} & \mathbf{100} & \mathbf{101} & \mathbf{110} & \mathbf{111} \\ - & 0 & 0 & 0 & 1 & 0 & 0 & 0 \\ 0 & - & 0 & 0 & 0 & 1 & 0 & 0 \\ 0 & 0 & - & 0 & 0 & 0 & 1 & 0 \\ 0 & 0 & 0 & - & 0 & 0 & 0 & 1 \\ 1 & 0 & 0 & 0 & - & 0 & 1 & 0 \\ 0 & 1 & 0 & 0 & 0 & - & 0 & 1 \\ 0 & 0 & 1 & 0 & 1 & 0 & - & 1 \\ 0 & 0 & 0 & 1 & 0 & 1 & 1 & - \end{pmatrix}$$

#### S9. Amalgamation of alternative rate matrices as independently evolving entities

In this section, I amalgamate the rate matrices, for all alternative coding schemes, as independently evolving (i.e., via SMM-ind) in order to analyze their behavior. The alternative coding schemes along with their scorings are summarized in Table S1 and Figure 4.

##### Scheme #1.

There are two characters: ***T*** - tail presence {*absent (a), present (p)*}, and ***C*** - tail color {*red (r), blue (b)*} that are defined by:

$$\boldsymbol{T} = \begin{pmatrix} \mathbf{0} & \mathbf{1} \\ -\alpha & \alpha \\ \beta & -\beta \end{pmatrix}, \ \boldsymbol{C} = \begin{pmatrix} \mathbf{0} & \mathbf{1} \\ -\gamma & \gamma \\ \theta & -\theta \end{pmatrix}, \text{(S9.1)}$$

Their amalgamation gives:

$$A_{mal.(ind)}(\boldsymbol{T},\boldsymbol{C}) = \boldsymbol{T}\otimes\boldsymbol{I_C} + \boldsymbol{I_T}\otimes\boldsymbol{C} = \begin{pmatrix} \mathbf{00} & \mathbf{01} & \mathbf{10} & \mathbf{11} \\ -\alpha-\gamma & \gamma & \alpha & 0 \\ \theta & -\alpha-\theta & 0 & \alpha \\ \beta & 0 & -\beta-\gamma & \gamma \\ 0 & \beta & \theta & -\beta-\theta \end{pmatrix}. \text{(S9.2)}$$

#### Scheme #2.

There are two characters: **T** - tail presence {*absent (0), present (1)*}, and **C** - tail color {no color (0), *red (1), blue (2)*}, defined by:

$$\boldsymbol{T}_2 = \begin{pmatrix} \mathbf{0} & \mathbf{1} \\ -\alpha & \alpha \\ \beta & -\beta \end{pmatrix}, \boldsymbol{C}_2 = \begin{pmatrix} \mathbf{0} & \mathbf{1} & \mathbf{2} \\ -\gamma-\delta & \gamma & \delta \\ \epsilon & -\epsilon-\eta & \eta \\ \theta & \lambda & -\theta-\lambda \end{pmatrix}. \text{ (S9.3)}$$

Their amalgamation gives:

$$A_{mal.(ind)}(\boldsymbol{T}_2, \boldsymbol{C}_2) = \begin{pmatrix} \mathbf{00} & \mathbf{01} & \mathbf{02} & \mathbf{10} & \mathbf{11} & \mathbf{12} \\ -\alpha-\gamma-\delta & \gamma & \delta & \alpha & 0 & 0 \\ \epsilon & -\alpha-\epsilon-\eta & \eta & 0 & \alpha & 0 \\ \theta & \lambda & -\alpha-\theta-\lambda & 0 & 0 & \alpha \\ \beta & 0 & 0 & -\beta-\gamma-\delta & \gamma & \delta \\ 0 & \beta & 0 & \epsilon & -\beta-\epsilon-\eta & \eta \\ 0 & 0 & \beta & \theta & \lambda & -\beta-\theta-\lambda \end{pmatrix}. \text{ (S9.4)}$$

#### Scheme #3.

There are three characters: $\boldsymbol{C_1}$ - tail presence {*absent (0), present (1)*}, $\boldsymbol{C_2}$ – red tail presence {*absent (0), present (1)*}, $\boldsymbol{C_3}$ – blue tail presence {*absent (0), present (1)*}:

$$\boldsymbol{C}_{3,1} = \begin{pmatrix} \mathbf{0} & \mathbf{1} \\ -\alpha & \alpha \\ \beta & -\beta \end{pmatrix}, \boldsymbol{C}_{3,2} = \begin{pmatrix} \mathbf{0} & \mathbf{1} \\ -\gamma & \gamma \\ \theta & -\theta \end{pmatrix}, \boldsymbol{C}_{3,3} = \begin{pmatrix} \mathbf{0} & \mathbf{1} \\ -\eta & \eta \\ \lambda & -\lambda \end{pmatrix}. \text{ (S9.5)}$$

$$A_{mal.(ind)}(\boldsymbol{C}_{3,1}, \boldsymbol{C}_{3,2}, \boldsymbol{C}_{3,3}) = \\ = \begin{pmatrix} \mathbf{000} & \mathbf{001} & \mathbf{010} & \mathbf{011} & \mathbf{100} & \mathbf{101} & \mathbf{110} & \mathbf{111} \\ -\alpha-\gamma-\eta & \eta & \gamma & 0 & \alpha & 0 & 0 & 0 \\ \lambda & -\alpha-\gamma-\lambda & 0 & \gamma & 0 & \alpha & 0 & 0 \\ \theta & 0 & -\alpha-\eta-\theta & \eta & 0 & 0 & \alpha & 0 \\ 0 & \theta & \lambda & -\alpha-\theta-\lambda & 0 & 0 & 0 & \alpha \\ \beta & 0 & 0 & 0 & -\beta-\gamma-\eta & \eta & \gamma & 0 \\ 0 & \beta & 0 & 0 & \lambda & -\beta-\gamma-\lambda & 0 & \gamma \\ 0 & 0 & \beta & 0 & \theta & 0 & -\beta-\eta-\theta & \eta \\ 0 & 0 & 0 & \beta & 0 & \theta & \lambda & -\beta-\theta-\lambda \end{pmatrix}. (S9.6)$$

#### Scheme #4.

This scheme is coded using one character: (1) tail - {*tail absent (0), tail red present (1), tail blue present (2)*}, and hence is represented by a single transition matrix by default:

$$\boldsymbol{T}_4 = \begin{pmatrix} \mathbf{0} & \mathbf{1} & \mathbf{2} \\ -\gamma-\delta & \gamma & \delta \\ \epsilon & -\epsilon-\eta & \eta \\ \theta & \lambda & -\theta-\lambda \end{pmatrix}. \text{ (S9.7)}$$

### S10. Amalgamation of alternative rate matrices as dependently evolving entities

In this section, I amalgamate the coding schemes by taking into account their specific dependencies; this converts all four coding schemes to SMM-ind and SMM-sw derived for the scheme #1 (see the equations S9.2 and S10.1).

#### Scheme #1.

The equation (S9.2) amalgamates the matrices from (S9.1) via SMM-ind. Alternatively, those matrices can be amalgamated through "switch-on" type of dependencies that gives:

$$A_{mal.(sw)}(\boldsymbol{T},\boldsymbol{C}) = \boldsymbol{T}\otimes\boldsymbol{I_C} + \boldsymbol{D_T}\otimes\boldsymbol{C} = \begin{pmatrix} \mathbf{00} & \mathbf{01} & \mathbf{10} & \mathbf{11} \\ -\alpha & 0 & \alpha & 0 \\ 0 & -\alpha & 0 & \alpha \\ \beta & 0 & -\beta-\gamma & \gamma \\ 0 & \beta & \theta & -\beta-\theta \end{pmatrix}. \text{ (S10.1)}$$

So, the scheme #1 can be represented by either SMM-ind (S9.2) or SMM-sw (S10.1).

#### Scheme #2.

Consider the amalgamated matrix from the equation (S9.4). The states 00 and 10 are improperly amalgamated using SMM-ind: the initial states within 00 (tail absent, no color) must change synchronously (since tail absence triggers immediate switch to no color); the initial states in the state 10 (tail present, no color) are illogical as tail must be always colored. So, these two amalgamated states, without loss of generality, have to be taken out from the matrix in the equation (S9.4) that produces the following rate matrix:

$$\boldsymbol{S_2} = \begin{pmatrix} \mathbf{01} & \mathbf{02} & \mathbf{11} & \mathbf{12} \\ -\alpha-\eta & \eta & \alpha & 0 \\ \lambda & -\alpha-\lambda & 0 & \alpha \\ \beta & 0 & -\beta-\eta & \eta \\ 0 & \beta & \lambda & -\beta-\lambda \end{pmatrix}.$$

We can notice that this matrix represents a matrix of two coevolving characters and hence can be aggregated with the respect to the two partitioning schemes {{01,02}, {11,12}} and {{01,11}, {02,12}} into two independent characters:

$$\boldsymbol{T} = \begin{pmatrix} \frac{1}{2} & 0 \\ 0 & \frac{1}{2} \end{pmatrix}\begin{pmatrix} 1 & 1 & 0 & 0 \\ 0 & 0 & 1 & 1 \end{pmatrix}\boldsymbol{S_2}\begin{pmatrix} 1 & 1 & 0 & 0 \\ 0 & 0 & 1 & 1 \end{pmatrix}^T = \begin{pmatrix} \mathbf{0} & \mathbf{1} \\ -\alpha & \alpha \\ \beta & -\beta \end{pmatrix},$$

$$\boldsymbol{C} = \begin{pmatrix} \frac{1}{2} & 0 \\ 0 & \frac{1}{2} \end{pmatrix}\begin{pmatrix} 1 & 0 & 1 & 0 \\ 0 & 1 & 0 & 1 \end{pmatrix}\boldsymbol{S_2}\begin{pmatrix} 1 & 0 & 1 & 0 \\ 0 & 1 & 0 & 1 \end{pmatrix}^T = \begin{pmatrix} \mathbf{1} & \mathbf{2} \\ -\eta & \eta \\ \lambda & -\lambda \end{pmatrix}.$$

These two characters are the same as tail presence (***T***) and tail color (***C***) characters from the coding scheme #1. So, the techniques used earlier for the scheme #1 can be applied to yield SMM-ind and SMM-sw from the equations S9.2 and S10.1.

#### Scheme #3.

Consider characters from the matrices (S3.5). The characters $\boldsymbol{C_{3,2}}$ (presence of red tail) and character $\boldsymbol{C_{3,3}}$ (presence of blue tail) change synchronously. Thus, they have to be amalgamated via SMM-syn that gives the character for the tail color {*red (0) or blue (1)*}:

$$A_{mal.(syn)}(\boldsymbol{C_{3,2}},\boldsymbol{C_{3,3}}) = \begin{pmatrix} \mathbf{0} & \mathbf{1} \\ -\gamma & \gamma \\ \theta & -\theta \end{pmatrix}.$$

The characters $A_{mal.(syn)}$ $(\boldsymbol{C}_{3,2}, \boldsymbol{C}_{3,3})$ and $\boldsymbol{C}_{3,2}$ are the same as those used in the scheme #1. Thus, the amalgamation techniques used for the scheme #1 can be applied to yield SMM-ind and SMM-sw from the equations S9.2 and S10.1.

#### S4a. Scheme #4.

The three-state matrix for the coding scheme #4 (matrix $\boldsymbol{T_4}$ in [S3.7]) cannot be further reduced; however, for comparison, its state space can be expanded using the lumpability property. This expansion implies constructing a four-state matrix that if lumped gives matrix $\boldsymbol{T_4}$. The convenient way to perform this expansion is to split the state *0* into the two states $0_1$ and $0_2$:

$$\boldsymbol{T}_{4ex} = \begin{pmatrix} \mathbf{0_0} & \mathbf{0_1} & \mathbf{1} & \mathbf{2} \\ -\gamma-\delta & 0 & \gamma & \delta \\ 0 & -\gamma-\delta & \gamma & \delta \\ \frac{\epsilon}{2} & \frac{\epsilon}{2} & -\epsilon-\eta & \eta \\ \frac{\theta}{2} & \frac{\theta}{2} & \lambda & -\theta-\lambda \end{pmatrix}.$$

To verify lumpability, the states in $\boldsymbol{T_{4ex}}$ have to be aggregated using the partitioning scheme {{*1,2*},{*3*},{*4*}}, this gives exactly $\boldsymbol{T_4}$:

$$\boldsymbol{T}_4 = \begin{pmatrix} \frac{1}{2} & 0 & 0 \\ 0 & 1 & 0 \\ 0 & 0 & 1 \end{pmatrix} \begin{pmatrix} 1 & 1 & 0 & 0 \\ 0 & 0 & 1 & 0 \\ 0 & 0 & 0 & 1 \end{pmatrix} \boldsymbol{T}_{4ex} \begin{pmatrix} 1 & 1 & 0 & 0 \\ 0 & 0 & 1 & 0 \\ 0 & 0 & 0 & 1 \end{pmatrix}^T = \begin{pmatrix} \mathbf{0} & \mathbf{1} & \mathbf{2} \\ -\gamma-\delta & \gamma & \delta \\ \epsilon & -\epsilon-\eta & \eta \\ \theta & \lambda & -\theta-\lambda \end{pmatrix}.$$

The matrix $\boldsymbol{T_{4ex}}$ can be further elaborated to model anatomical dependencies. For example, to construct SMM-sw of the scheme #1 the left diagonal elements of $\boldsymbol{T_{4ex}}$ have to be set to zeros. The SMM-ind of the scheme #1 can be obtained from the previous example by relaxing the assumption of no transitions for the "*absent*" states.

#### S11. Two-scientist paradox

Even though the characters in the simulations evolve independently, let us consider their joint evolution by amalgamating the shape ***S*** {*round ($r_o$), triangular (t)*} and color ***C*** {*blue (b), red (r)*} characters:

$$A_{mal.(ind)}(\boldsymbol{S},\boldsymbol{C}) = \boldsymbol{S}\otimes\boldsymbol{I_C} + \boldsymbol{I_S}\otimes\boldsymbol{C} = \begin{pmatrix} \boldsymbol{r_o b} & \boldsymbol{r_o}\mathbf{r} & \boldsymbol{tb} & \boldsymbol{tr} \\ -\alpha-\gamma & \gamma & \alpha & 0 \\ \theta & -\alpha-\theta & 0 & \alpha \\ \beta & 0 & -\beta-\gamma & \gamma \\ 0 & \beta & \theta & -\beta-\theta \end{pmatrix} \text{(S11.1).}$$

This matrix is the same with the one used in the simulation for the two-scientist paradox; it has four free parameters. The aggregation of states under the discretization schemes used by the first scientists $\mathcal{B}_{1.1}$={{ $r_o$*b*, $r_o$*r* }, {*tb,tr*}}, $\mathcal{B}_{1.2}$={{$r_o$*b*, *tb* }, {$r_o$*r*, *tr*}}, $\mathcal{B}_{1.3}$={{$r_o$*b*}, {*tb* }, {$r_o$*r*}, {*tr*}} always produces lumpable models that correspond to the characters: shape, color and shape + color respectively. The aggregation used by the second scientist $\mathcal{B}_{2.1}$={{$r_o$*b*, $r_o$*r tbl*}, {*tr*}} violates the row-wise sum rule, and hence the original model is not lumpable under this scheme. If all rates in the matrix S11.1 are equal (i.e., the matrix is defined by one free parameter), then the row-wise sum rule does not hold for the discretization scheme of the second scientist as well.

*Simulations.* One hundred trees of 2000 taxa were simulated using *phytools* package (Revell 2012). This number of taxa was used to demonstrate the asymptotic behavior of models under conditions when the amount of data is sufficient. The two-state color and shape characters were simulated as independent on the generated trees. To simulate different rates, the first rate of the shape character was drawn from a uniform distribution [0, 0.1], then this value was multiplied by the factors 1.5 and 0.1 to yield the second rate as well as the rates of the color character; so each character was represented by the two unique rates. To create the scoring for the first scientist, the characters were left intact. For the second scientist, the states of the simulated characters were recoded to produce a two-state character: {*red triangular present, red triangular absent*}. Next, both HMM and MM were fitted for each character (except the first scientist's simultaneous character of shape + color since its original characters were simulated as independent given the initial conditions) using *rayDisc* function of *corHMM* package.
