## Supplementary material for "Integration of anatomy ontologies and evo-devo using structured Markov models suggests a new framework for modeling discrete phenotypic traits": SMM_examples.R

```
##############################################################################################
#
# EXAMPLES OF CONSTRUCTING STRUCTURED MARKOV MODELS (SMM, SMM+HMM)
# FOR TREE INFERENCE AND COMPARATIVE ANALYSIS
#
# Sergei Tarasov 7/06/2018
#
##############################################################################################

# set working directory
setwd("/Scripts_to_construct_SMMs")

# use functions from this file
source("SMM_functions.R")

#####################
# TAIL COLOR PROBLEM (GENERAL MODEL)
#####################
# initilize tail character
char.state<-c("a", "p")
rate.param<-c(1, 2)
TL<-init_char_matrix(char.state, rate.param, diag.as=0)
diag(TL)<-rate.param*-1
TL
# initilize color character
char.state<-c("r", "b")
rate.param<-c(3, 4)
COL<-init_char_matrix(char.state, rate.param, diag.as=0)
diag(COL)<-rate.param*-1
COL
# amalgamating characters as independently evolving (SMM-ind)
TC.ind<-comb2matrices(TL, COL, controlling.state=NULL, name.sep="", diag.as="")
TC.ind

### amalgamating characters via swithed-on dependency (SMM-sw)
TC.sw<-comb2matrices(TL, COL, controlling.state=2, name.sep="", diag.as="")
TC.sw

#####################
# TAIL ARMOR CASE (GENERAL MODEL)
#####################
# initilize tail character
char.state<-c("Ta", "Tp")
rate.param<-c(1, 2)
TL<-init_char_matrix(char.state, rate.param, diag.as=0)
diag(TL)<-rate.param*-1
TL

# armor character
char.state<-c("Aa", "Ap")
rate.param<-c(3, 4)
AR<-init_char_matrix(char.state, rate.param, diag.as=0)
diag(AR)<-rate.param*-1
AR

# armor color
char.state<-c("R", "B")
rate.param<-c(5, 6)
COL<-init_char_matrix(char.state, rate.param, diag.as=0)
diag(COL)<-rate.param*-1
COL

# amalgamating characters as independently evolving (SMM-ind)
ARM.ind<-comb2matrices(TL, comb2matrices(AR, COL, controlling.state=NULL, name.sep="", diag.as=0),
                       controlling.state=NULL, name.sep="", diag.as=0)
ARM.ind

######################################################
#
# Mk-SMMs FOR REVBAYES
#
#######################################################
# copy the outputs generated by cat(...) and paste in Tree_inference_SMM.Rev

# since all parameters are the same in Mk-like
# models, let's construct a new amalgamated character accordingly.
# use fnFreeK(...rescaled=true...) in RevBayes to rescale the matrix properly

###################################
# SMMs for the tail color problem
###################################
char.state<-c("a", "p")
rate.param<-c(1, 1)
TL<-init_char_matrix(char.state, rate.param, diag.as=0)
char.state<-c("r", "b")
rate.param<-c(1, 1)
COL<-init_char_matrix(char.state, rate.param, diag.as=0)

#SMM-ind
TC.ind<-comb2matrices(TL, COL, controlling.state=NULL, name.sep="", diag.as="")
TC.ind
in.rev<-Mk_Rev(TC.ind)
cat(in.rev) # COPY the output and insert in Rev template Tree_inference_SMM.Rev
cat(in.rev, file="input_Rev.txt") # or save this outout to a file and then copy to Rev template

#SMM-sw
TC.sw<-comb2matrices(TL, COL, controlling.state=2, name.sep="", diag.as="")
TC.sw
cat(Mk_Rev(TC.sw)) # COPY the output and insert in Rev template Tree_inference_SMM.Rev

###################################
# SMMs for the tail armor case
###################################
# initilize tail character
char.state<-c("Ta", "Tp")
rate.param<-c(1, 1)
TL<-init_char_matrix(char.state, rate.param, diag.as=0)
TL
# armor character
char.state<-c("Aa", "Ap")
rate.param<-c(1, 1)
AR<-init_char_matrix(char.state, rate.param, diag.as=0)
AR
# armor color
char.state<-c("R", "B")
rate.param<-c(1, 1)
COL<-init_char_matrix(char.state, rate.param, diag.as=0)
COL

# SMM-sw
ARM.sw<-comb2matrices(AR, COL, controlling.state=2, name.sep="", diag.as="")
ARM.sw<-comb2matrices(TL, ARM.sw, controlling.state=2, name.sep="", diag.as="")
cat(Mk_Rev(ARM.sw))

# SMM-ind
ARM.ind<-comb2matrices(AR, COL, controlling.state=NULL, name.sep="", diag.as="")
ARM.ind<-comb2matrices(TL, ARM.ind, controlling.state=NULL, name.sep="", diag.as="")
cat(Mk_Rev(ARM.ind))

########################################################
#
# SMM-sw FOR corHMM
# (the provided ouputs can be used with other packages
#   if adjusted accordingly)
#
#####################################################

# matrix to simulate
char.state<-c("a", "p")
rate.param<-c(0.1, 0.2)
TL<-init_char_matrix(char.state, rate.param, diag.as=0)
diag(TL)<-rate.param*-1

char.state<-c("r", "b")
rate.param<-c(0.05, 0.15)
COL<-init_char_matrix(char.state, rate.param, diag.as=0)
diag(COL)<-rate.param*-1

Q.sw<-comb2matrices(TL, COL, controlling.state=2, name.sep="", diag.as="")
Q.sw

# simulate history
require("phytools")
tree<-pbtree(n=100, scale=200)
sim.h<-sim.history(tree, Q.sw, nsim=1, anc=setNames(c(1, 0, 0, 0), colnames(Q.sw) ) )
plot(sim.h)

# matrix for inference
char.state<-c("a", "p")
rate.param<-c(1, 2)
TL<-init_char_matrix(char.state, rate.param, diag.as=0)

char.state<-c("r", "b")
rate.param<-c(3, 4)
COL<-init_char_matrix(char.state, rate.param, diag.as=0)

SMM.sw<-comb2matrices(TL, COL, controlling.state=2, name.sep="", diag.as="", non.rate.as=NA)
SMM.sw

# recode simulated character by aggregating ar and ab states; so the final character
# will have only three states {a, pr, pb}, and the state (a) will consiste of the two hidden states
require("plyr")
char.recode<-mapvalues(sim.h$states, from = c("ar", "ab"), to = c("ar&ab", "ar&ab"))


# reconstruct ancestral states using corHMM
# WARNING: in this exmple I use literal state names (e.g., ab..), if it does not work use numbers instead 
require("corHMM")
taxa=cbind(sim.h$tip.label, char.recode)
out.smm <- rayDISC(sim.h, taxa, rate.mat=SMM.sw, node.states="marginal", model="ARD", root.p="maddfitz")
```
