## Supplementary material for "Integration of anatomy ontologies and evo-devo using structured Markov models suggests a new framework for modeling discrete phenotypic traits": SMM_functions.R

```
##############################################################################################
# FUNCTIONS
##############################################################################################

#' @title Combining two matrices
#' @description Combining two matrices. The parametric schem of matrice is defined by nattural
#' numbers; different numbers = different rate parameters
#' @param M1 matrix; if dependency true thenM1 controls M2
#' @param M2 matrix; if dependency true then: M2 depends on those states of M1 specified in controlling.state
#' @param controlling.state state(s) of M1 that switches on/off matrix M2 
#' @param name.sep separator for state names
#' @param diag.as hpopulate main diagonal with
#' @return Matrix
#' @examples
#' M1<-matrix(c(-1,1,  2,-2),2,2,byrow=TRUE)
#' rownames(M1)<-colnames(M1)<-c("0","1")
#' M2<-matrix(c(-3,3,  4,-4),2,2,byrow=TRUE)
#' rownames(M2)<-colnames(M2)<-c("0","1")
#' comb2matrices(M1, M2, controlling.state=NULL)
#' comb2matrices(M1, M2, controlling.state=2)

comb2matrices<-function(M1,M2, controlling.state=NULL, name.sep="", diag.as="", non.rate.as=NULL){
  
  if (!is.null(controlling.state)){
    matrix.diag<-rep(0, ncol(M1))
    matrix.diag[controlling.state]<-1
    matrix.diag<-diag(matrix.diag)
  }
  
  if (is.null(controlling.state)){
    matrix.diag<-diag(nrow(M1))
  }
  
  M_kr=(M1%x%diag(nrow(M2)))+ (matrix.diag%x%M2)
  
  
  #getting colnames
  
  col=paste(colnames(kronecker(M1, diag(nrow(M2)), make.dimnames = T)),
            colnames(kronecker(diag(nrow(M1)), M2, make.dimnames = T)), sep="")
  col=gsub("::", name.sep, col, fixed=T)
  
  # merging two names
  rownames(M_kr)<-colnames(M_kr)<-col
  if (diag.as!="") diag(M_kr)<-diag.as
  if (!is.null(non.rate.as)) M_kr[which(M_kr<=0)]<-non.rate.as
  
  return(M_kr)
}


#' @title Initialize binary matrices
#' @param char.state names for character states
#' @param rate.param names for the rate parameters
#' @param diag.as values to pas to the main diagonal elements
#' @return matrix

init_char_matrix<-function(char.state, rate.param, diag.as=NA){
  n.state<-length(char.state)
  Q=matrix(ncol = n.state, nrow=n.state,byrow=TRUE)
  Q[xor(lower.tri(Q, diag = FALSE), upper.tri(Q, diag = FALSE))]<-rate.param
  Q<-t(Q)
  diag(Q)<-diag.as
  rownames(Q)<-colnames(Q)<-as.character(char.state)
  return(Q)
}

#' @title Make rate matrix for RevBayes
#' @param M rate matrix
#' @param eq.diag keep main diagonal elements equal and rescale off-diagonal elements
#' @return matrix
#' @examples
#' cat(Mk_Rev(M))

Mk_Rev<-function(M, eq.diag=FALSE){
  
  dec<-paste("
NUM_STATES=", nrow(M), "\n",
             "for (i in 1:NUM_STATES) {
  for (j in 1:NUM_STATES) {
      rates[i][j] <-0.0
  }
}\n", sep="", collapse="")
  
  rt<-c()
  for (i in 1:nrow(M)){
    which(M[i,]>0)->IJ
    
    rates<-M[i,IJ]
    
    if (eq.diag==TRUE) rates<-M[i,IJ]/sum(M[i,IJ])
    
    # convert integer to decimal (RevBAyes sometimes retuns error if decimals are not used)
    rates[rates%%1==0]<-paste(rates[rates%%1==0], ".0", sep="")
    
    rt<-c(rt,
          paste("rates[", i, "][", IJ, "]<-", rates, "\n", sep ="" )
    )}
  
  rt<-paste(rt, collapse="")
  rt<-paste(dec, rt, sep = "")
  
  return(rt)
}
#########
```
