## Supplementary material for "Integration of anatomy ontologies and evo-devo using structured Markov models suggests a new framework for modeling discrete phenotypic traits": Sim_2ScieParadox_pipiline.R

```
########
# FUNCTIONS

# simulate under random matrices
k0=1.5
k1=0.1
#k2=0.2
nsim=100

LL<-sim_hist_rnd(k0=1.5, k1=0.1,  nsim=100)

sim_hist_rnd<-function(k0, k1,  nsim){
  
  LL<-list(M=vector("list", nsim), simA=vector("list", nsim), simB=vector("list", nsim)  )
  
  for (i in 1:nsim){
  tree<-pbtree(n=2000, scale=200)
  r12<-runif(1, 0, 0.1)  
  # LL$M[[i]]$QA<-QA<-RM(char.state=c(1,2), rate.param=c(r12, r12*k0))
  # LL$M[[i]]$QB<-QB<-RM(char.state=c(1,2), rate.param=c(r12*k1, r12*k0*k1))
  LL$M[[i]]$QA<-QA<-RM(char.state=c(1,2), rate.param=c(r12, r12*k0))
  LL$M[[i]]$QB<-QB<-RM(char.state=c(1,2), rate.param=c(r12*k0*k1, r12*k1))
  
  LL$simA[[i]]<-simA<-sim.history(tree, QA, nsim=1, anc=setNames(c(1, 0), c(1,2)))
  LL$simB[[i]]<-simB<-sim.history(tree, QB, nsim=1, anc=setNames(c(0, 1), c(1,2)))
  }
  
return(LL)  
}
LL$simA
LL$M
LL$simA[[1]]$
# tree<-pbtree(n=2000, scale=200)
# 
# QA<-RM(char.state=c(1,2), rate.param=c(0.05, 0.03))
# QB<-RM(char.state=c(1,2), rate.param=c(0.005, 0.007))
# 
# 
# simA<-sim.history(tree, QA, nsim=10, anc=setNames(c(1, 0), c(1,2)))
# simB<-sim.history(tree, QB, nsim=10, anc=setNames(c(0, 1), c(1,2)))

###
# chack distribution of state combination per run
hid.tb<-mapply(
  function(x, y){
    S<-paste(x$states,  y$states, sep = ":")
    unique(S)
    table(S)
  },
  x=LL$simB, y=LL$simA
)
######
#####
# make hidden states
S.hid<-mapply(
  function(x, y){
    S<-paste(x$states,  y$states, sep = ":")
    mapvalues(S, from = c("1:1", "1:2", "2:1", "2:2"), to = c("1&2&3", "1&2&3", "1&2&3", "4"))
  },
  x=LL$simB, y=LL$simA
)

S.hidA<-mapply(
  function(x, y){
    S<-paste(x$states,  y$states, sep = ":")
    mapvalues(S, from = c("1:1", "1:2", "2:1", "2:2"), to = c("1&2&3", "1&2&3", "1&2&3", "4"))
  },
  x=LL$simB, y=LL$simA
)


cbind(LL$simB[[1]]$states,
LL$simA[[1]]$states)
S.hid[,1]
[1] "1&2&3" "1&2&3" "1&2&3" "1&2&3" "1&2&3" "1&2&3" "1&2&3" "1&2&3" "1&2&3" "1&2&3" "4"  
######
# make two states
S.lump<-mapply(
  function(x, y){
    S<-paste(x$states,  y$states, sep = ":")
    mapvalues(S, from = c("1:1", "1:2", "2:1", "2:2"), to = c(1, 1, 1, 2))
  },
  x=LL$simB, y=LL$simA
)
######
# hidden matrix
# hidden matrix
char.state<-c(1, 2)
rate.param<-c(1, 2) # rate parameters = (n.states^2)-n
QA.rr<-init_char_matrix(char.state, rate.param, diag.as=0)
rate.param<-c(3, 4) # rate parameters = (n.states^2)-n
QB.rr<-init_char_matrix(char.state, rate.param, diag.as=0)

Q.hid<-comb2matrices(QA.rr, QB.rr, dependent.state=NULL)
Q.hid[Q.hid==0]<-NA
rownames(Q.hid)<-colnames(Q.hid)<-c(1:4)
##############
#######################
# reconstruct HIDDEN
######################
LL$simA[[1]]

i=1
recon.hid<-vector("list", nsim)
for (i in 1:nsim){
taxa=cbind(LL$simA[[i]]$tip.label, S.hid[,i])
tree.recon=LL$simA[[i]]
rate.mat=Q.hid
recon.hid[[i]] <- rayDISC(tree.recon, taxa, rate.mat=rate.mat, node.states="marginal", model="ARD", root.p="maddfitz")
}

###############################
## non hidden embeded in hidden (rates equally distributed)
#
##################
# matrix
char.state<-c(1, 2, 3, 4)
rate.param<-c(NA) # rate parameters = (n.states^2)-n
QA.E<-init_char_matrix(char.state, rate.param, diag.as=NA)
QA.E[1:3,]<-1
QA.E[4,]<-2
diag(QA.E)<-NA
###
recon.hid.E<-vector("list", nsim)
for (i in 1:nsim){
taxa=cbind(LL$simA[[i]]$tip.label, S.hid[,i])
tree.recon=LL$simA[[i]]
rate.mat=QA.E
recon.hid.E[[i]] <- rayDISC(tree.recon, taxa, rate.mat=rate.mat, node.states="marginal", model="ARD", root.p="maddfitz")
}

############################
## no hidde, 2-rates
#
##################
# matrix
char.state<-c(1, 2)
rate.param<-c(1, 2) # rate parameters = (n.states^2)-n
QA.r<-init_char_matrix(char.state, rate.param, diag.as=NA)
###
recon.2r<-vector("list", nsim)
for (i in 1:nsim){
  taxa=cbind(LL$simA[[i]]$tip.label, S.lump[,i])
  tree.recon=LL$simA[[i]]
  rate.mat=QA.r
  recon.2r[[i]] <- rayDISC(tree.recon, taxa, rate.mat=rate.mat, node.states="marginal", model="ARD", root.p="maddfitz")
}

########################################################################################
#
# Hidden vs nonhidden for states for original blue and red characters
#
########################################################################################

############################
## no hidde, 2-rates
#
##################
# matrix A
char.state<-c(1, 2)
rate.param<-c(1, 2) # rate parameters = (n.states^2)-n
QA.r<-init_char_matrix(char.state, rate.param, diag.as=NA)
###
i=1
LL$simA[[1]]$states
simA.2r<-vector("list", nsim)
for (i in 1:nsim){
  taxa=cbind(LL$simA[[i]]$tip.label, LL$simA[[i]]$states)
  tree.recon=LL$simA[[i]]
  rate.mat=QA.r
  simA.2r[[i]] <- rayDISC(tree.recon, taxa, rate.mat=rate.mat, node.states="marginal", model="ARD", root.p="maddfitz")
}

##################
# matrix B
char.state<-c(1, 2)
rate.param<-c(1, 2) # rate parameters = (n.states^2)-n
QA.r<-init_char_matrix(char.state, rate.param, diag.as=NA)
###
i=1
LL$simB[[1]]$states
simB.2r<-vector("list", nsim)
for (i in 1:nsim){
  taxa=cbind(LL$simB[[i]]$tip.label, LL$simB[[i]]$states)
  tree.recon=LL$simB[[i]]
  rate.mat=QA.r
  simB.2r[[i]] <- rayDISC(tree.recon, taxa, rate.mat=rate.mat, node.states="marginal", model="ARD", root.p="maddfitz")
}
############################################
######
# hidden matrix
char.state<-c(1, 2)
rate.param<-c(1, 2) # rate parameters = (n.states^2)-n
QA.rr<-init_char_matrix(char.state, rate.param, diag.as=0)
rate.param<-c(3, 4) # rate parameters = (n.states^2)-n
QB.rr<-init_char_matrix(char.state, rate.param, diag.as=0)

Q.hid<-comb2matrices(QA.rr, QB.rr, dependent.state=NULL)
Q.hid[Q.hid==0]<-NA
rownames(Q.hid)<-colnames(Q.hid)<-c(1:4)
#
Q.hid[2,4]<-2
Q.hid[3,1]<-1
#
Q.hid[1,2]<-1
Q.hid[2,1]<-1
Q.hid[2,4]<-1

Q.hid[3,1]<-2
Q.hid[3,4]<-2
Q.hid[4,3]<-2
##############
# expand state space
LL$simA[[1]]$states

mapvalues(LL$simA[[1]]$states, from = c("1", "2"), to = c("1&2", "3&4"))
simA.exp<-lapply(LL$simA, function(x) mapvalues(x$states, from = c("1", "2"), to = c("1&2", "3&4")))
simB.exp<-lapply(LL$simB, function(x) mapvalues(x$states, from = c("1", "2"), to = c("1&2", "3&4")))
#
#######################
# reconstruct HIDDEN
######################
LL$simA[[1]]

i=1
simA.hid<-vector("list", nsim)
for (i in 1:nsim){
  taxa=cbind(LL$simA[[i]]$tip.label, simA.exp[[i]])
  tree.recon=LL$simA[[i]]
  rate.mat=Q.hid
  simA.hid[[i]] <- rayDISC(tree.recon, taxa, rate.mat=rate.mat, node.states="marginal", model="ARD", root.p="maddfitz")
}
#####
LL$simB[[1]]

i=1
simB.hid<-vector("list", nsim)
for (i in 1:nsim){
  taxa=cbind(LL$simB[[i]]$tip.label, simB.exp[[i]])
  tree.recon=LL$simB[[i]]
  rate.mat=Q.hid
  simB.hid[[i]] <- rayDISC(tree.recon, taxa, rate.mat=rate.mat, node.states="marginal", model="ARD", root.p="maddfitz")
}
############
simA.2r[[100]]$AIC-simA.hid[[100]]$AIC
simA.2r[[100]]$AIC

a.aic.m<-lapply(simA.2r, function(x) x$AIC) %>% unlist
a.aic.hm<-lapply(simA.hid, function(x) x$AIC)%>% unlist
hist(a.aic.m-a.aic.hm)
min(a.aic.hm-a.aic.m)
max(a.aic.hm-a.aic.m)
mean(a.aic.hm-a.aic.m)
max(a.aic.m-a.aic.hm)

b.aic.m<-lapply(simB.2r, function(x) x$AIC) %>% unlist
b.aic.hm<-lapply(simB.hid, function(x) x$AIC)%>% unlist
b.aic.m-b.aic.hm
hist(b.aic.m-b.aic.hm)
max(b.aic.m-b.aic.hm)

min(b.aic.hm-b.aic.m)
max(b.aic.hm-b.aic.m)
min(b.aic.hm-b.aic.m)

#sep.chars=list(simA.2r=simA.2r, simA.hid=simA.hid, simB.2r=simB.2r, simB.hid=simB.hid)
#saveRDS(sep.chars, file="sep.chars_shape_col.rds")


h3A<-lapply(hid3$simA.hid, function(x) x$AIC)%>% unlist
h2A<-lapply(hid2$simA.hid, function(x) x$AIC)%>% unlist
h1A<-lapply(sep.chars$simA.hid, function(x) x$AIC)%>% unlist

hist(h3A-h2A)
max(h3A-h2A)
mean(h3A-h2A)
hist(h3A-h1A)
mean(h3A-h1A)
max(h3A-h1A)

Fn<-ecdf(h3A-h1A)
Fn(10)
Fn(-10)
Fn(-4)
hid3$simA.hid[[1]]
sep.chars$simA.hid[[1]]

h3B<-lapply(hid3$simB.hid, function(x) x$AIC)%>% unlist
h2B<-lapply(hid2$simB.hid, function(x) x$AIC)%>% unlist
h1B<-lapply(sep.chars$simB.hid, function(x) x$AIC)%>% unlist

hist(h3B-h2B)
mean(h3B-h2B)
max(h3B-h2B)
Fn<-ecdf(h3B-h1B)
Fn(-10)

hist(h3B-h1B)
mean(h3B-h1B)
max(h3B-h1B)

daic<-seq(-15, 15, 0.1)
plot(daic, -1*(4+daic)/2, type="l", col="green", lwd=2)
abline(v=0)
abline(v=-10)
abline(v=10)
abline(h=3)
abline(h=-7)
abline(h=0)

-1*(4+10)/2
###########################
#Hid 2
# > Q.hid
# 1  2  3  4
# 1 NA  3  1 NA
# 2  4 NA NA  2
# 3  1 NA NA  3
# 4 NA  2  4 NA

#hid2<-list(simA.hid=simA.hid, simB.hid=simB.hid)
#saveRDS(hid2, file="hid2_shape_col.rds")

###########################
#Hid 3
# > Q.hid
# 1  2  3  4
# 1 NA  1  1 NA
# 2  1 NA NA  1
# 3  2 NA NA  2
# 4 NA  2  2 NA

#hid3<-list(simA.hid=simA.hid, simB.hid=simB.hid)
#saveRDS(hid3, file="hid3_eg_shape_col.rds")
################
# save
#saveRDS(recon.hid, file="recon_hid.rds")
#saveRDS(recon.hid.E, file="recon_hid_E.rds")
#saveRDS(recon.2r, file="recon_2r.rds")
#

# compare rate estimation in non-hidden models
######## 
rE12<-lapply(recon.hid.E, function(x) x$solution[1,4]) %>% unlist
rE21<-lapply(recon.hid.E, function(x) sum(x$solution[4,c(1:3)]) ) %>% unlist

eq12<-lapply(recon.2r, function(x) x$solution[1,2]) %>% unlist
eq21<-lapply(recon.2r, function(x) x$solution[2,1]) %>% unlist

h12<-lapply(recon.hid, function(x) mean(x$solution[c(2:3),4]) ) %>% unlist
h21<-lapply(recon.hid, function(x) sum(x$solution[4, c(2:3)]) ) %>% unlist

##inital states used in simulation
comb2matrices(QA.rr, QB.rr, dependent.state=NULL)
cbind(LL$simB[[1]]$states,
      LL$simA[[1]]$states)

# delta sim-Hid
(comb2matrices(LL$M[[1]]$QB, LL$M[[1]]$QA, dependent.state=NULL)[c(2, 3, 5, 9)]-
  recon.hid[[1]]$solution[c(2, 3, 5, 9)] ) %>% abs %>% sum

(comb2matrices(LL$M[[1]]$QA, LL$M[[1]]$QB, dependent.state=NULL)[c(2, 3, 5, 9)]-
  recon.hid[[1]]$solution[c(2, 3, 5, 9)] ) %>% abs %>% sum
#
simBA<-lmapply(LL$M, function(x) comb2matrices(x$QB,x$QA, dependent.state=NULL)[c(2, 3, 5, 9)]  )
simAB<-lapply(LL$M, function(x) comb2matrices(x$QA,x$QB, dependent.state=NULL)[c(2, 3, 5, 9)]  )

simH<-lapply(recon.hid, function(x) x$solution[c(2, 3, 5, 9)])

plot(unlist(simAB), unlist(simH))
plot(lapply(simAB, sum) %>% unlist, lapply(simH, sum) %>% unlist)
# error
dABH<-mapply(function(x, y){
  abs(x-y) %>% sum },
  x=simAB, y=simH
)

dABHs<-mapply(function(x, y){
 abs(sum(x[1:2])-sum(y[1:2]) )^2 +
  abs((sum(x[3:4])/2-sum(y[3:4])/2))^2
 },
  x=simAB, y=simH
)

dBAH<-mapply(function(x, y){
  abs(x-y) %>% sum },
  x=simBA, y=simH
)
mean(dBAH)
mean(dABH)

mean(dABHs)

dABH<dBAH
hist(dABH)
sd(dABH)
sd(dABHs)

# delta sim - nonHid
simABS<-lapply(simAB, function(x) c(sum(x[c(3,4)])/2, sum(x[c(1,2)] ) ) )
NH<-lapply(recon.2r, function(x) c(x$solution[1,2], x$solution[2,1]) )

dABNH<-mapply(function(x, y){
  abs(x-y)^2 %>% sum },
  x=simABS, y=NH
)
mean(dABNH)
hist(dABNH)
sd(dABNH)
plot(unlist(simABS), unlist(NH))
recon.2r[[1]]$solution
  
####
plot(rE12, eq12)
plot(rE21, eq21)
plot(h12, eq12)
plot(h21, eq21)

LL$M[[1]]
recon.hid.E[[1]]$solution
recon.2r[[1]]$solution
recon.hid[[1]]$solution
recon.hid.E[[1]]$loglik
recon.2r[[1]]$loglik

#########

recon.hid[[1]]$AIC
del<-(lapply(recon.hid.E, function(x) x$AIC) %>% unlist)-
  (lapply(recon.hid, function(x) x$AIC) %>% unlist)

del.e<-(lapply(recon.hid.E, function(x) x$AIC) %>% unlist)-
  (lapply(recon.2r, function(x) x$AIC) %>% unlist)
max(del.e)
min(del.e)

hist(del, breaks = 10)
min(del)
which(del==min(del))
Fn<-ecdf(del)
Fn(10)
Fn(10)
Fn(2)
summary(Fn)

plot(Fn)
plot(Fn, verticals = TRUE, do.points = FALSE)

plot(del,
hid.tb[4,]
)
LL$M[[3]]
lapply(LL$M, function(x) x$QA[1,2]) %>% unlist %>% plot(del,.)
lapply(LL$M, function(x) x$QA[1,2]) %>% unlist->r1

# regression
hid.tb
reg<-as_tibble(cbind(del, t(hid.tb)/2000, r1) )
colnames(reg)<-c("del", "c11", "c12", "c21", "c22", "r1" )
lm<-lm(del ~ c11, data=reg)
lm<-lm(del ~ c12+r1, data=reg)
lm<-lm(del ~ poly(c12,3)+r1, data=reg)
summary(lm)
AIC(lm)
plot(lm)
plot(reg$c12^3, reg$del)
plot( reg$r1, reg$del)

del.cut<-cut(del, c(-Inf,0, 10,Inf))
plot(reg$r1, reg$c12, col=del.cut)
scatter3D(reg$r1, reg$c12, reg$del)
plotrgl()

hist(del,  breaks=30,freq = TRUE, include.lowest = F, right = F)
h<-hist(del,  breaks=20, plot=FALSE, probability = TRUE, right = F)
cuts <- cut(h$breaks, c(-Inf,0,10,Inf))
cuts<-mapvalues(cuts, from = c("(-Inf,0]", "(0,10]", "(10, Inf]"), to = brewer.pal(9, 'Set1')[1:3])
cuts<-c("black", brewer.pal(9, 'Set1')[9], brewer.pal(9, 'Set1')[4])[c(cuts)]
par(  mar=c(5,5,5,5))
plot(h, col=cuts,  main="", xlab=expression(paste(Delta, "AIC", sep="")), xlim = c(-5, 80), xaxt = "n",  yaxt = "n", cex.lab=2)
axis(side = 1, at=c(-5, 5, 15, 40, 80), label=c(-5, 0, 10, 40, 80),cex.axis=1.5)
axis(side = 2, at=seq(0, 12, 4), label=seq(0, 12, 4)/100, cex.axis=1.5)
#abline(h=10, v=10)
```
